# Increased epithelial mTORC1 activity in chronic rhinosinusitis with nasal polyps

**DOI:** 10.1101/2023.10.13.562288

**Authors:** George X. Huang, Nils R. Hallen, Minkyu Lee, Kelly Zheng, Xin Wang, Michael V. Mandanas, Sarah Djeddi, Daniela Fernandez, Jonathan Hacker, Tessa Ryan, Regan W. Bergmark, Neil Bhattacharyya, Stella Lee, Alice Z. Maxfield, Rachel E. Roditi, Kathleen M. Buchheit, Tanya M. Laidlaw, James E. Gern, Teal S. Hallstrand, Anuradha Ray, Sally E. Wenzel, Joshua A. Boyce, Maria Gutierrez-Arcelus, Nora A. Barrett

**Affiliations:** Jeff and Penny Vinik Center for Translational Immunology Research, Division of Allergy and Clinical Immunology, Brigham and Women’s Hospital; Boston, MA; Department of Medicine, Harvard Medical School; Boston, MA; Department of Immunology, Harvard Medical School; Boston, MA; Division of Immunology, Boston Children’s Hospital; Boston, MA; Department of Otolaryngology, Head and Neck Surgery, Brigham and Women’s Hospital; Boston, MA; Department of Otolaryngology, Head and Neck Surgery, Massachusetts Eye and Ear Infirmary; Boston, MA; Program in Medical and Population Genetics, Broad Institute of MIT and Harvard; Cambridge, MA; Division of Allergy, Immunology, and Rheumatology, University of Wisconsin School of Medicine and Public Health; Madison, WI; Division of Pulmonary, Critical Care and Sleep Medicine, University of Washington Medical Center; Seattle, WA; Department of Immunology, University of Pittsburgh; Pittsburgh, PA; Department of Pulmonary, Allergy and Critical Care Medicine, University of Pittsburgh Medical Center; Pittsburgh, PA

**Keywords:** epithelial cell, basal cell, airway stem cell, basal cell adhesion molecule, airway inflammation, type 2 inflammation, interleukin-4, interleukin-13, chronic rhinosinusitis with nasal polyps, asthma, bulk RNA-sequencing, single cell RNA-sequencing

## Abstract

**Background:** The airway epithelium plays a central role in the pathogenesis of chronic respiratory diseases such as asthma and chronic rhinosinusitis with nasal polyps (CRSwNP), but the mechanisms by which airway epithelial cells (EpCs) maintain inflammation are poorly understood.

**Objective:** We hypothesized that transcriptomic assessment of sorted airway EpCs across the spectrum of differentiation would allow us to define mechanisms by which EpCs perpetuate airway inflammation.

**Methods:** Ethmoid sinus EpCs from adult patients with CRS were sorted into 3 subsets, bulk RNA sequenced, and analyzed for differentially expressed genes and pathways. Single cell RNA-seq (scRNA-seq) datasets from eosinophilic and non-eosinophilic CRSwNP and bulk RNA-seq of EpCs from mild/moderate and severe asthma were assessed. Immunofluorescent staining and *ex vivo* functional analysis of sinus EpCs were used to validate our findings.

**Results:** Analysis within and across purified EpC subsets revealed an enrichment in glycolytic programming in CRSwNP vs CRSsNP. Correlation analysis identified mammalian target of rapamycin complex 1 (mTORC1) as a potential regulator of the glycolytic program and identified EpC expression of cytokines and wound healing genes as potential sequelae. mTORC1 activity was upregulated in CRSwNP, and *ex vivo* inhibition demonstrated that mTOR is critical for EpC generation of CXCL8, IL-33, and CXCL2. Across patient samples, the degree of glycolytic activity was associated with T2 inflammation in CRSwNP, and with both T2 and non-T2 inflammation in severe asthma.

**Conclusions:** Together, these findings highlight a metabolic axis required to support epithelial generation of cytokines critical to both chronic T2 and non-T2 inflammation in CRSwNP and asthma.

**KEY MESSAGES:**

- Epithelial mTORC1 activity is upregulated in CRSwNP.
- mTOR regulates EpC cytokine generation.
- Epithelial metabolic reprograming correlates with T2 inflammation in CRSwNP, and with both T2 and non-T2 inflammation in asthma.

**CAPSULE SUMMARY:** mTORC1 mediates EpC cytokine generation in CRSwNP.

## INTRODUCTION

Genomic and transcriptomic studies have identified a central role for the airway epithelium in the pathogenesis of inflammatory respiratory diseases such as asthma and chronic rhinosinusitis with nasal polyps (CRSwNP).^1–4^ While the advent of therapeutic monoclonal antibodies that target signaling via IL-4Rα^5,6^ and IL-5^7,89^ has been a major advance for the treatment of CRSwNP and the common T2^high^ endotype of asthma,^10^ there is a need to better define the mechanisms by which the airway epithelium perpetuates and sustains T2 and non-T2 inflammation in chronic airway diseases.

Several studies using single cell RNA sequencing (scRNA-seq) of epithelial cells (EpCs) in CRSwNP have identified top alterations in highly expressed genes that reflect the epithelial response to T2 cytokines. Our group reported the first study using scRNA-seq in CRS,^11^ finding that basal EpCs from CRSwNP expressed a robust set of T2 cytokine-induced genes including *ALOX15* (which encodes arachidonate 15-lipoxygenase), *POSTN*, *PTHLH*, *SERPINB2*, and *CCL26*. In another scRNA-seq study, Stevens et al. found increased expression of *ALOX15* in sinonasal EpCs from subjects with aspirin-exacerbated respiratory disease (AERD), a variant of CRSwNP characterized by severe T2 inflammation, as compared to aspirin-tolerant controls.^12^ Moreover, *ALOX15* expression was correlated with nasal eosinophilic cationic protein, reflecting eosinophil tissue burden.^12^ A scRNA-seq study by Kotas et al. featuring >100,000 EpCs confirmed that expression of *ALOX15* and *POSTN* are induced by IL-13 in airway EpCs,^13^ while a more recent scRNA-seq study including healthy controls, CRS without nasal polyps (CRSsNP), non-eosinophilic CRSwNP (neCRSwNP), and eosinophilic CRSwNP (eCRSwNP) identified that overexpression of *ALOX15* is also detected in a population of type 2 conventional dendritic cells (cDC2s) that is uniquely present in eCRSwNP.^14^ Genomic studies have identified variants in *ALOX15* as risk factors for CRSwNP development,^4^ and mechanistic studies have demonstrated that *ALOX15* leads to glutathione depletion^15^ and oxidized phosphatidyl ethanolamine metabolites that lead to ferroptosis,^16,17^ and also promotes production of *CCL26* (eotaxin-3) in airway EpCs.^18^ Thus, T2 cytokine-driven upregulation of epithelial *ALOX15* is a likely feed-forward pathway by which EpCs maintain T2 inflammation.

Another observation that has emerged from scRNA-seq studies of CRSwNP is the prominence of cellular/tissue remodeling, often associated with the response to T2 cytokines. We determined that patients with CRSwNP have basal cell hyperplasia, wherein basal cells fail to differentiate normally,^11^ but rather persist in a stem-like state that can be maintained through IL-4/13 and insulin receptor substrate (IRS) signaling.^19^ A recent scRNA-seq study identified an increased tuft cell frequency in CRSwNP, as compared to healthy control mucosa.^13^ Here investigators used a murine model to show that chronic exposure to IL-13 (one month duration) was sufficient to expand tracheal tuft cell numbers. Notably, while goblet cell hyperplasia is also identified in this disease, it is not unique to the T2^high^ phenotype, but is also present in CRSsNP.^20^ Moreover, no study in CRSwNP has detected the mucociliary EpC state identified in T2 inflammation in asthma,^1^ suggesting interesting differences between epithelial response to T2 cytokines in the upper and lower airways. Finally, scRNA-seq studies and histologic assessments have reliably identified a reduction in submucosal gland cells in CRSwNP,^11,21^ which is poorly understood.

Taken together, the recent literature has emphasized alterations in lipid mediator biology and cellular remodeling as dominant changes in the airway epithelium of CRSwNP, often associated with T2 inflammation Airway EpCs exhibit great plasticity and are highly tuned to their surrounding milieu.^22^ Accordingly, airway EpC gene expression can predict individuals with T2 inflammation and therapeutic glucocorticoid responsiveness.^10^ However, individual reports using the power of scRNA-seq to assess EpC gene expression are constrained by small sample sizes and the sparsity of data,^11,19^ while larger studies using bulk RNA-seq are limited by the inability to detect individual cell states. Thus, we have not yet fully leveraged the potential of transcriptomic studies in the respiratory epithelium to define EpC dysfunction and endotype airway diseases.

We recently reported that human airway basal cells express high levels of basal cell adhesion molecule (BCAM), which reliably distinguishes this progenitor population from differentiating EpCs.^19^ In this study, we sorted highly purified BCAM^hi^ basal cells, transitional EpCs, and differentiated EpCs from subjects with CRS and performed bulk RNA-seq to define novel mechanisms that support epithelial inflammation. Here we identify that CRSwNP is distinguished by prominent mTORC1 activity and that mTORC regulates the generation of select EpC cytokines. Furthermore, we show that while mTORC1-dependent genes can be driven by both T2 and T17 cytokines *in vitro*, mTORC1-related metabolic reprogramming correlates with the degree of T2 inflammation and wound healing in CRSwNP and correlates with the degree of both T2 and T17 inflammation in asthma.

## MATERIALS AND METHODS

### Bulk RNA-seq analysis of human sorted EpC subsets

Ethmoid sinus tissue was collected from human subjects between the ages of 18 and 75 who underwent endoscopic sinus surgery at Brigham and Women’s Hospital (Boston, MA) for CRSwNP (n=11) or chronic rhinosinusitis without nasal polyps (CRSsNP) (n=8) (**Table E1**). The Mass General Brigham Institutional Review Board approved the study and all subjects provided written informed consent prior to participation.

Specimens were chopped, digested in RPMI-1640 medium (ThermoFisher 11875093) with 10% FBS containing type IV collagenase (Worthington LS004189) and DNaseI (Sigma 10104159001) with magnetic stir bar at 600 RPM at 37°C for 30 min, and triturated using a 25 mL syringe with 16g needle every 15 min. The resultant cell suspension was filtered through a 70μm cell strainer, centrifuged at 500g for 10min, washed, and resuspended. Single cell suspensions were sorted using the gating strategy shown in **Fig 1A**. In brief, cells were blocked with Fc receptor blocker (BioLegend 101320) for 10 min on ice, then incubated with fluorophore-labeled antibodies to human EpCAM (BioLegend 118213), NGFR (Abcam ab52987), BCAM (MBL International D295-3), CD31 (BioLegend 2434), CD45 (BioLegend 103116), and CD90 (BioLegend 105328) for 30 min.

**Fig 1:**
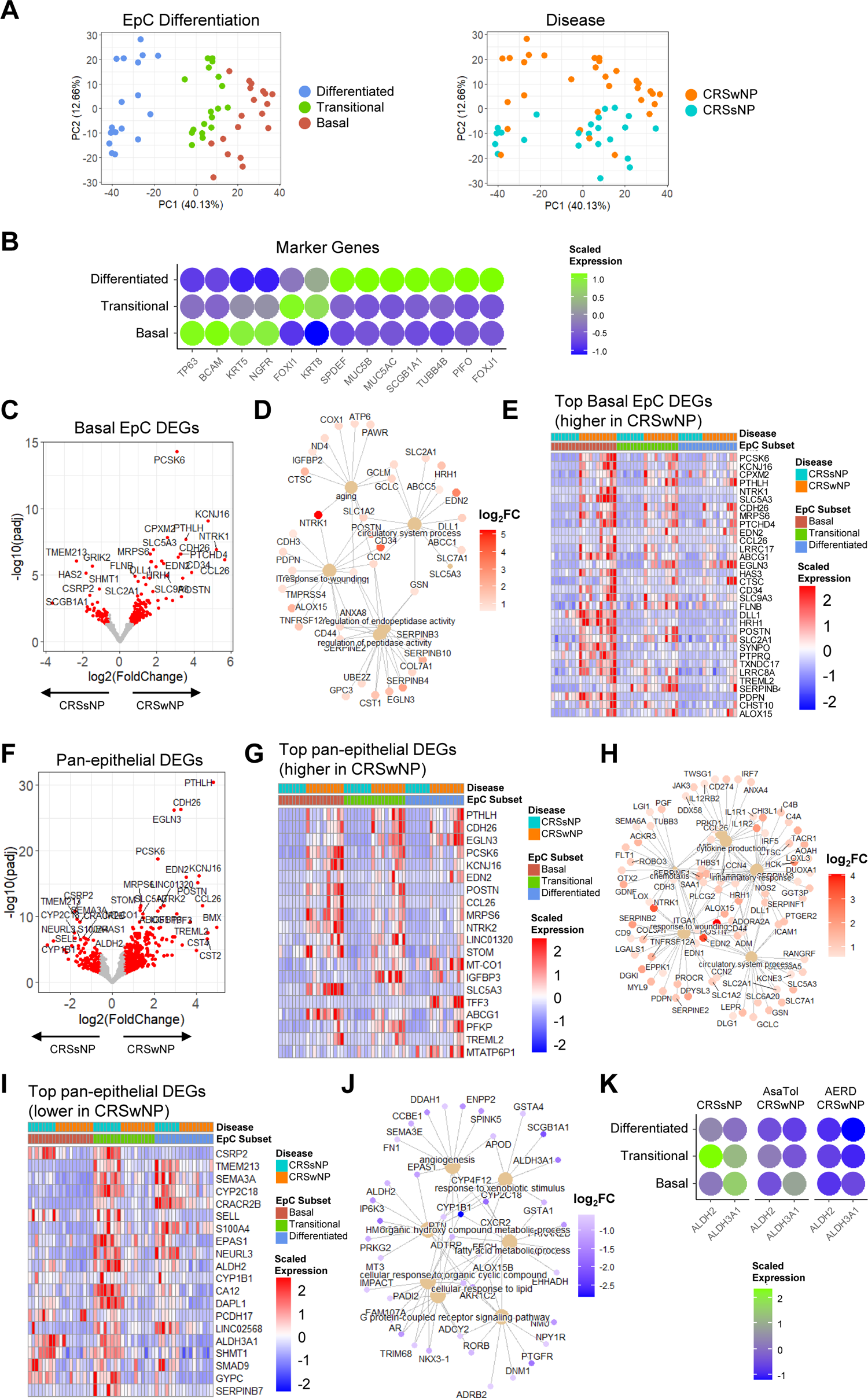
EpCs from CRSwNP are enriched for wound healing and metabolic genes and have lost key protective factors. A) PCA plots (PC1 and PC2) of bulk-RNA seq samples. Left is colored by EpC subset. Right is colored by disease. B) Dot plot of canonical EpC markers for basal, transitional, and differentiated EpCs. C) Volcano plot of CRSwNP vs CRSsNP DE testing in basal EpCs. Positive log_2_FoldChange indicates higher expression in CRSwNP. Red indicates DEGs meeting |log_2_FoldChange|>0.58 and padj<0.05. D) Net plot of selected top over-represented gene sets (in the gene ontology biological processes database) for DEGs with higher expression in CRSwNP basal EpCs. E) Heatmap of top DEGs with higher expression in CRSwNP basal EpCs. F) Volcano plot of pan-epithelial CRSwNP vs CRSsNP DE testing. Positive log_2_FoldChange indicates higher expression in CRSwNP. Red indicates DEGs meeting |log_2_FoldChange|>0.58 and padj<0.05. G) Heatmap of top pan-epithelial DEGs with higher expression in CRSwNP EpCs. H) Net plot of selected top over-represented gene sets (in the gene ontology biological processes database) for pan-epithelial DEGs with higher expression in CRSwNP EpCs. I) Heatmap of top pan-epithelial DEGs with lower expression in CRSwNP EpCs. J) Net plot of selected top over-represented gene sets (in the gene ontology biological processes database) for pan-epithelial DEGs with lower expression in CRSwNP EpCs. K) Dot plot of *ALDH2* and *ALDH3A1* by cell subset and disease (including aspirin tolerant CRSwNP and AERD).

7AAD (BioLegend 420403) was added immediately before cell sorting. After removing debris, doublets, and dead cells, EpCAM^+^ Lineage^-^ EpCs were sorted into three populations: BCAM^high^ EpCAM^low^ NGFR^+^ basal EpCs, EpCAM^high^ NGFR^+^ transitional EpCs, and EpCAM^high^ NGFR^-^ differentiated EpCs. 1000 cells were collected for each population, mixed with 5ul TCL buffer (Qiagen 1031576), and stored at −80°C. RNA libraries were prepared using Smart-Seq2 and 38bp paired-end sequencing was performed by the Harvard-MIT Broad Institute.

Sequencing quality was assessed with FastQC.^23^ Reads were pseudo-aligned and transcript expression quantified using Kallisto^24^ with the Ensembl GRCh38 transcriptome. Quantification files were then processed for downstream analysis with the Tximport pipeline.^25^ Samples were considered acceptable for downstream analysis if they had >90% of commonly expressed genes (defined as genes detected in 80% of samples) and Pearson’s *r*>0.5 with all other samples. To enhance discovery and reduce computational demand, genes that did not meet a minimum detection threshold within the dataset (raw count>12 in at least 15% of samples) were removed. PCA using the top 500 most variable genes (after variance stabilizing transformation of size factor-normalized counts)^26^ confirmed samples separated by EpC and there were no outliers (**Fig 1B**). Gene expression was normalized using DESeq2’s median of ratios method,^27^ and differential expression testing was performed within DESeq2 while including sex, oral corticosteroid usage, and pseudoalignment rate as covariates. P-values were adjusted for multiple testing using the Benjamini-Hochberg method.^28^ Genes meeting both |log_2_FoldChange|>0.58 (which corresponds to a base 10 foldchange of ±50%) and padj<0.05 were considered to be differentially expressed (DE).

Over-representation analysis (ORA) and gene set enrichment analysis (GSEA)^29^ were performed using the clusterProfiler package^30^ with the human MSigDb gene ontology (GO) biological processes and Hallmark databases,^31,32^ respectively. For GSEA of the Hallmark glycolysis gene set, the gene IL13RA1 was excluded to prevent bias. Gene set variation analysis (GSVA) was used to score samples for expression of selected gene sets,^33^ and the Limma package^34^ was used to remove possible effects from oral corticosteroid usage prior to generating normalized counts for GSVA. Of note, the IL-13 response score was only calculated for 10 of 11 subjects with CRSwNP, as one study subject had received dupilumab (IL-4Rα antagonist) before undergoing sinus surgery. Spearman correlation calculation was performed between GSVA scores and other scores or gene expression data (ρ indicates Spearman’s *rho*). When applicable for correlation between GSVA scores, any overlapping genes were excluded from the score on the y-axis (for example, overlapping genes in the Hallmark glycolysis and mTORC1 signaling scores were excluded from the mTORC1 signaling score in **Fig 2D** and **Fig 4B**).

**Fig 2:**
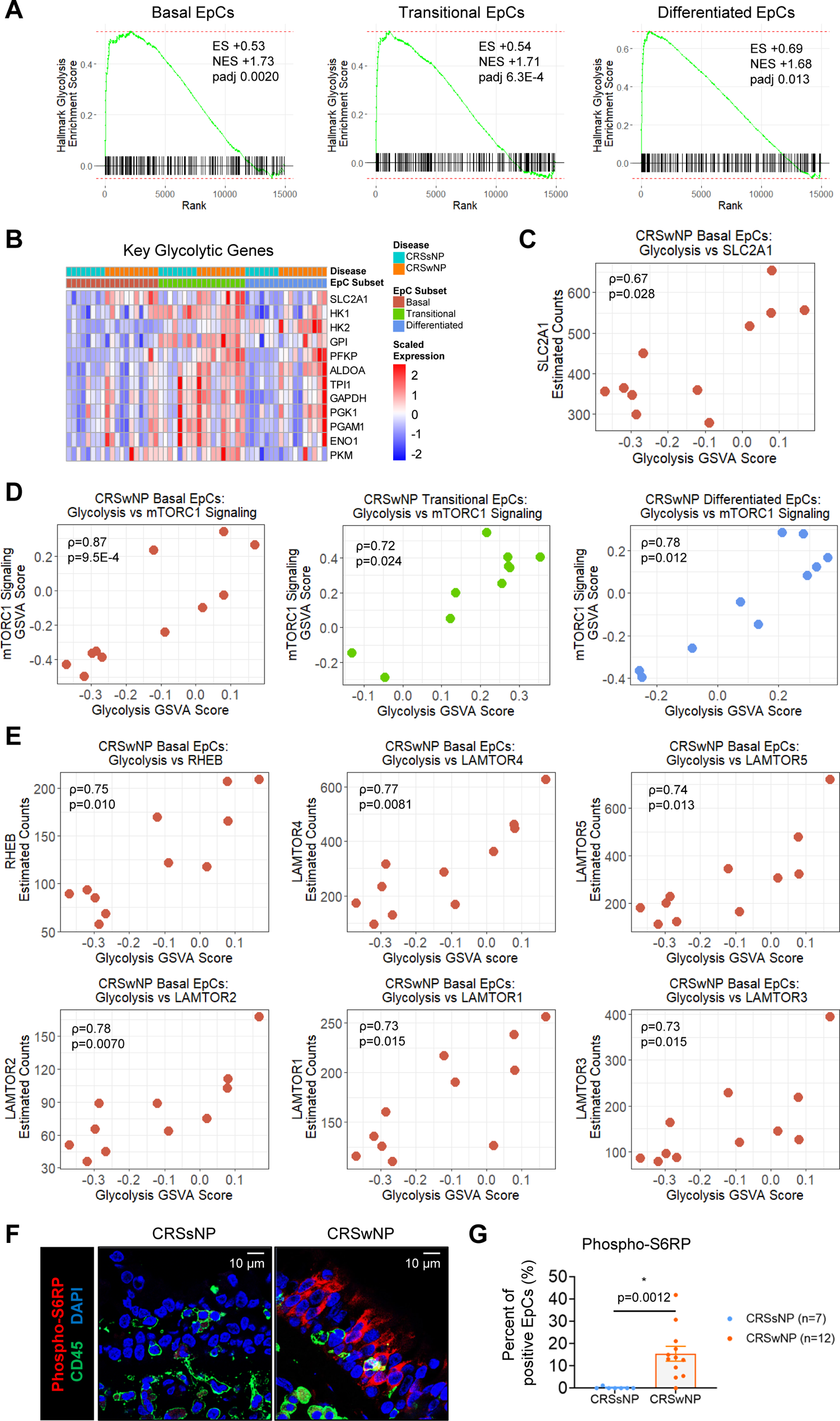
The enhanced mTORC1 signaling in CRSwNP EpCs is tightly correlated with glycolysis. A) GSEA plots of the Hallmark glycolysis gene set for CRSwNP vs CRSsNP DE testing in basal EpCs, transitional EpCs, and differentiated EpCs. Positive ES or NES indicates enrichment in CRSwNP. *IL13RA1* was excluded from the glycolysis gene set to prevent bias. B) Heatmap of key genes in the glycolysis pathway. C) Scatterplot of glycolysis GSVA score vs *SLC2A1* estimated counts in CRSwNP basal EpCs. *ρ* indicates Spearman’s *rho*. D) Scatterplots of glycolysis GSVA score vs mTORC1 signaling GSVA score in CRSwNP basal EpCs, CRSwNP transitional EpCs, and CRSwNP differentiated EpCs. E) Scatterplots of glycolysis GSVA score vs estimated counts of mTORC1 regulators (*RHEB*, *LAMTOR4*, *LAMTOR5*, *LAMTOR2*, *<u>LAMTOR1</u>*, *LAMTOR3*) in CRSwNP basal EpCs. F) Representative images of phospho-S6RP, CD45, and DAPI staining in ethmoid sinus samples from adult human donors with CRSsNP and CRSsNP. G) Quantification of percentage of phospho-S6RP^positive^ cells out of CD45^negative^ cells in the epithelial layer of adult human donors with CRSsNP and CRSsNP. The non-parametric Mann-Whitney U test was used for statistical testing.

### Bulk RNA-seq analysis of the publicly available bronchial ALI dataset

Bulk RNA-seq FASTQ files and metadata from human bronchial epithelial cell (HBEC) air-liquid interface (ALI) cultures stimulated with a variety of cytokines were downloaded from the NCBI Gene Expression Omnibus [GSE185202],^22^ and pseudo-aligned using Kallisto^24^ as above. Differential expression testing was performed with DESeq2,^27^ using donor as a covariate. GSEA was performed using clusterProfiler^30^ and the Hallmark gene set database. The top 200 differentially expressed genes induced by each cytokine (the 200 genes with lowest padj and positive log_2_FoldChange in response to cytokine stimulation) were used to construct epithelial cytokine response signatures for IL-13, IL-17, IFN-α, and IFN-γ (**Table E10**); these cytokine signatures were used to score samples in other datasets using GSVA^33^ (**Fig 6E**, **Fig 6F**) or Seurat module scoring.^35,36^ Genes were considered IL-13 responsive (**Fig E7**) if they exhibited log_2_FoldChange>0 and padj<0.05 in response to IL-13 stimulation.

### Single cell RNA-seq (scRNA-seq) analysis of the publicly available Kotas dataset

A Seurat object of sinus EpCs from healthy control (n=4) and CRSwNP (n=5) was downloaded from the NCBI Gene Expression Omnibus [GSE202100].^13^ The original cell-type annotations and SCT gene expression values were used for visualization and analysis. Module scores were calculated using Seurat’s built-in AddModuleScore function.^35,36^

### scRNA-seq (scRNA-seq) analysis of the publicly available Wang dataset

Metadata and scRNA-seq FASTQ files containing epithelial and immune cells from healthy control (n=5), CRSsNP (n=5), non-eosinophilic CRSwNP (neCRSwNP) (n=5), and eosinophilic CRSwNP (eCRSwNP) (n=6) were downloaded from the Genome Sequence Archive [HRA000772]^14^ and aligned to Gencode GRCh38 using 10x Genomics Cell Ranger v6.1.2 with default parameters.^37^ Cells with >12,000 UMIs, >15% mitochondrial reads, or <500 genes detected were filtered out, as they were considered to be either fragments or doublets. RNA counts were scaled using the NormalizeData function prior to downstream analysis or visualization. Principal component analysis was performed on top variable genes and PCs were subsequently corrected using the Harmony package^38^ to correct for disease (“Public_Description”) and sex (“Gender”).

UMAPs were generated using Harmonized principal components, and the dataset was sequentially re-clustered into various immune and epithelial populations consistent with the original report^14^ (**Fig E4** and **Tables E11-20**). Gene expression module scores were calculated using Seurat’s built-in AddModuleScore function.^35,36^ A cluster of proliferating cells (**Fig E4A**) was excluded prior to cell type quantification as this was comprised of mixed immune, epithelial, and stromal cells. A cluster of ciliated epithelial cells was also excluded prior to cell type quantification due to poor capture in the neCRSwNP and eCRSwNP nasal polyp samples.^14^

### Bulk RNA-seq analysis of the publicly available IMSA bronchial brushing dataset

Bulk RNA-seq FASTQ files and metadata for bronchial brushing samples (healthy control n=17, mild-to-moderate asthma n=25, severe asthma n=23) from the Immune Mechanisms of Severe Asthma (IMSA) bronchial brushing study were downloaded from the NCBI Gene Expression Omnibus [GSE158752],^39,40^ and pseudo-aligned using Kallisto^24^ as above. Differential expression testing was performed with DESeq2,^27^ using sex, age, and sequencing batch as covariates. Genes meeting both |log_2_FoldChange|>0.58 (which corresponds to a base 10 foldchange of ±50%) and padj<0.05 were considered to be differentially expressed (DE). GSEA was performed using clusterProfiler^30^ and the Hallmark gene set database.

### Immunofluorescence of human sinonasal tissue

Fresh human ethmoid sinus surgical samples from CRSwNP and CRSsNP were fixed overnight with 4% paraformaldehyde (Boston BioProducts BM-155) at 4°C, washed with HBSS for 20 min, and dehydrated in 30% sucrose overnight at 4°C. Tissue was then embedded in OCT compound (Fisher 23-730-571), cryosectioned at 5 μm thickness using a cryostat (Leica CM1850), and adhered to positively charged glass slides (Fisher 12-550-15). Cryosections were blocked with 1x blocking buffer (Abcam ab126587) in PBST with 0.2% Triton X-100 and 0.1% Tween-20 for 1 h at room temperature. Afterwards, slides were incubated with primary antibodies to human phospho-6SRP (Ser235/236) (CellSignaling 4858) or CD45 (BioLegend 304002) at 1:50 dilution overnight at 4°C, washed 3x with PBS, incubated with the appropriate secondary antibodies (Invitrogen AlexaFluor488 and AlexaFluor594 conjugates) at 1:200 dilution for 1 h at room temperature, washed 3x with PBS, and mounted with DAPI mounting medium (Abcam ab104139). Images were obtained with a Zeiss LSM 800 Laser Scanning Confocal Microscope. Images were analyzed and merged using ImageJ (National Institutes of Health, Bethesda, MD). Percent of phospho-6SRP^positive^ cells in the epithelial layer per high power field (HPF) was quantified out of the total CD45^negative^ cells in the epithelial layer per HPF.

### Immunocytochemistry of cultured CRSwNP basal EpCs

Human basal EpCs (BCAM^high^ EpCAM^low^ NGFR^+^) from subjects with CRSwNP were expanded in PneumaCult Ex basal medium (STEMCELL 05008) containing 500μL StemCell Hydrocortisone (STEMCELL 07925) and 1% Penicillin-Streptomycin (ThermoFisher 15140122). After 3 passages, basal EpCs were seeded onto 8-well chamber slides. Cells were stimulated with 20 ng/mL IL-1β (BioLegend 579402) and 20 ng/mL TNF-α (BioLegend 570104) for 48 h, in the presence or absence of 5 nM Torin 1 (Selleck Chem S2827). Cells were then fixed with 4% paraformaldehyde for 10 mins and incubated overnight with antibodies to human CXCL2 (LSBio B14609), CXCL8 (R&D MAB208), or IL-33 (R&D AF3625) using the recommended manufacturers’ dilutions, followed by the appropriate secondary antibodies. Slides were mounted using DAPI mounting medium (Abcam ab104139), and fluorescence images were obtained with a Zeiss LSM 800 Laser Scanning Confocal Microscope. Images were analyzed and merged using ImageJ^41^ (National Institutes of Health, Bethesda, MD). Normalized integrated density was quantified by taking the average integrated density divided by the number of nuclei per HPF.

### Statistical analyses

Statistical analyses were performed using GraphPad Prism v9, Seurat v4,^35,36^ and DESeq2 v3.17.^27^ R packages were implemented in RStudio^42^ with R version 4.1.2. Where applicable, non-parametric Mann-Whitney U tests or Wilcoxon rank-sum tests were used for statistical comparisons.

### Data availability

Bulk RNA-seq expression data will be available via NCBI GEO.

## RESULTS

### EpCs from CRSwNP are enriched for wound healing and metabolic genes and have lost key protective factors

Live, lineage negative (CD31^-^, CD45^-^, CD90^-^) EpCs from 19 human ethmoid sinus specimens (11 CRSwNP and 8 CRSsNP) were sorted into basal (EpCAM^low^, NGFR^+^, BCAM^high^), transitional (EpCAM^high^, NGFR^+^), and differentiated EpCs (EpCAM^high^, NGFR^-^) (**Fig E1**) and then subjected to bulk RNA-seq. Principal component analysis (PCA) using the top 500 most variable genes showed that PC1 (explaining 40.1% of variance) captured the spectrum of EpC differentiation, with basal EpC genes^11,19^ (e.g. *KRT5, TP63)* contributing strongly to positive scores on PC1, and with differentiated EpC genes^11,13^ (e.g. *SCGB1A1, MUC5AC)* contributing strongly to negative scores on PC1 (**Fig 1A**). PC2 (explaining 12.7% of variance) captured the spectrum of disease, with previously reported CRSwNP genes^11,13^ such as *PTHLH*, *POSTN*, and *ALOX15* contributing strongly to positive scores on PC2 (**Fig 1A**). As expected, basal EpCs demonstrated high expression of marker genes^11,19^ such as *KRT5, TP63, BCAM, and NGFR*, while differentiated EpCs had high expression of ciliated (*PIFO*, *FOXJ1*, *TUBB4B*) and secretory (*SCGB1A1*, *MUC5B*, *MUC5AC*) genes (**Fig 1B**). Interestingly, transitional EpCs had high expression of the ionocyte marker FOXI1 (**Fig 1B**).^43^

Differential expression (DE) testing for CRSwNP vs CRSsNP within basal EpCs, while controlling for sex and OCS usage as covariates, demonstrated 260 differentially expressed genes (DEGs) (|log_2_FoldChange|>0.58 and padj<0.05) for CRSwNP vs CRSsNP, including 188 DEGs with higher expression in CRSwNP basal EpCs (**Fig 1C, Table E2**). Among the top basal EpC DEGs with higher expression in CRSwNP were genes associated with airway T2 inflammation such as *PTHLH*, *CDH26*, *CCL26*, *POSTN*, and *ALOX15*.^11,44,45^ Additionally, two top DEGs *SLC9A3* and *SYNPO* were previously reported to be IL-13-inducible, upregulated in the esophagus of patients with active eosinophilic esophagitis, and important in barrier function,^46,47^ underscoring the potential for shared barrier dysfunction across these T2 disorders. Other top DEGs included those involved in maintenance of stemness or inhibition of EpC differentiation such as the notch ligand *DLL1,*^48^ and the hematopoietic stem cell marker *CD34*, now appreciated to maintain the stem potential of diverse stromal cell types.^49,50^ This is consistent with our recent work demonstrating an increased number of BCAM^hi^ basal cells in CRSwNP.^19^ Finally, basal cells from CRSwNP also had high levels of expression of genes involved in metabolism including *SLC5A3*, encoding a sodium and myo-inositol co-transporter that activates cell proliferation through an Akt/mTOR pathway,^51^ and *SLC2A1*, encoding the glucose transporter GLUT1 that is upstream of the glycolysis pathway.^52,53^ Over-representation analysis of the 188 DEGs with higher expression in CRSwNP basal EpCs yielded GO biological processes gene sets involved in wound healing (padj=0.028) and secreted peptidase inhibitors (padj=0.0061) (**Fig 1D**), perhaps consistent with DEGs involved in barrier function, stemness, and growth.

Visual inspection of the top basal EpC DEGs in CRSwNP compared to CRSsNP showed that many appeared to have preserved differential expression throughout the spectrum of EpC differentiation (**Fig 1E**). Moreover, formal DE testing between CRSwNP and CRSsNP within transitional (**Fig E1B, Table E3**) and differentiated EpCs (**Fig E1C, Table E4**) identified many of the same DEGs. Given the notable overlap in DEGs across EpC subsets, we enhanced our resolution by performing DE analysis for CRSwNP vs CRSsNP across all samples while controlling for EpC subset, sex, and OCS usage as covariates. We identified 359 pan-epithelial DEGs with higher expression in CRSwNP vs CRSsNP (log_2_FoldChange>0.58 and padj<0.05) and 137 pan-epithelial DEGs with lower expression in CRSwNP vs CRSsNP (log_2_FoldChange<-0.58 and padj<0.05) (**Fig 1F**). Among the top pan-epithelial DEGs with higher expression in CRSwNP were expected genes of T2 inflammation (including *PTHLH*, *CDH26*, *POSTN*, *CCL26*), as well as *PFKP* (log_2_FoldChange=1.29, padj=6.72E-10) which encodes a rate limiting enzyme of glycolysis,^54,55^ and *SLC2A1* (log_2_FoldChange=0.713, padj=1.82E-6) (**Fig 1G**). In addition to these differences in T2 and metabolic genes, we detected higher expression of genes involved in regulation of growth factor signaling (*IGFBP3*) and the hypoxia response (*EGLN3*, which encodes a prolyl hydroxylase that post-translationally modifies hypoxia-inducible factor 1α, HIF-1α).^56^ Over-representation analysis of the 359 pan-epithelial DEGs with higher expression in CRSwNP again revealed gene sets involved in wound healing, as well as a more prominent role for genes involved the inflammatory response (padj=4.4E-5), chemotaxis (padj=7.2E-5), and cytokine production (padj=0.031) (**Fig 1H**). Among the top pan-epithelial DEGs with lower expression in CRSwNP were genes involved in protection against oxidative stress such as *ALDH2* (log_2_FoldChange=-0.902, padj=4.25E-7) and *ALDH3A1* (log_2_FoldChange=-1.63, padj=2.78E-6) (**Fig 1I-K, Table E5**). Interestingly, we observed that *ALDH2* expression tended to be lower in CRSwNP subjects with AERD vs aspirin-tolerant CRSwNP (**Fig 1K**). Genetic polymorphisms causing ALDH2 deficiency contribute to ethanol-induced cutaneous and respiratory reactions^57,58^ as well as increased risk of various epithelial malignancies,^59^ and we speculate that the reduced *ALDH2* expression here may underly the alcohol intolerance that is commonly observed in patients with AERD.^60^

### Enhanced mTORC1 signaling in CRSwNP EpCs is tightly correlated with glycolysis

Gene set enrichment analysis (GSEA) using the Hallmark gene sets demonstrated that glycolysis was positively enriched in CRSwNP within each EpC subset (basal padj=0.0020, transitional padj=6.3E-4, differentiated padj=0.013) (**Fig 2A**) as well as in the pan-epithelial comparison (pan-epithelial padj=5.6E-4) (**Fig E2**). Additionally, similar to prior reports,^53^ higher expression of key glycolytic genes was also evident in CRSwNP vs CRSsNP across EpC subsets (**Fig 2B**). To better assess the importance of glycolytic programming across replicate patient samples, we used gene set variation analysis (GSVA) to score the transcriptome of individual CRSwNP samples for expression of Hallmark glycolysis genes. GSVA demonstrated a wide range of scores within this group (**Fig 2C**), so we leveraged this variation to identify co-expressed genes and potential molecular regulators. We observed that *SLC2A1* (GLUT1, ρ=0.67 and p=0.028), which is not a member of the Hallmark glycolysis gene set (**Fig 2C**), correlated with expression of the glycolysis score, consistent with a role for GLUT1 in regulating EpC glycolytic activity.^53^ We also observed that the glycolysis score was tightly correlated with mTORC1 signaling in all CRSwNP EpC subsets (basal ρ=0.87, p=9.5E-4; transitional ρ=0.72, p=0.024; differentiated ρ=0.78, p=0.012) (**Fig 2D**). Furthermore, focusing on the highly purified basal EpC population, we found significant correlations between glycolysis and positive regulators of mTORC1 signaling, including *RHEB*, *LAMTOR1*, and every member of the Ragulator complex (**Fig 2E).**^61,62^ In contrast, we did not detect statistically significant positive correlations between glycolysis and a PI3K-AKT-MTOR gene signature, which is more representative of mTORC2 signaling (**Fig E3**).^63^ We validated epithelial mTORC1 activity at the protein level across 19 additional subjects with CRS, demonstrating robust phosphorylated S6 ribosomal protein (phospho-S6RP) in CRSwNP that was absent in CRSsNP (**Fig 2F**, **Fig 2G)**.

As mTORC1 signaling regulates a plethora of downstream pathways, we sought to clarify which mTORC1-dependent genes may be responsible for enhancement of glycolysis in CRSwNP EpCs. Among the genes in the Hallmark mTORC1 signaling gene set, those which were most positively correlated with the basal EpC glycolysis score were *CD9* (ρ=0.92, p=6.7E-5) and *SHMT2* (ρ=0.85, p=8.1E-4), as well as genes with known glycolytic function including *GAPDH* (ρ=0.91, p=1.1E-4) and *PGM1* (ρ=0.92, p=6.7E-5). *CD9* encodes a membrane-spanning protein that has previously been reported to enhance glycolytic activity in tumor cells^64^ and has been implicated as an inducer of cellular senescence.^65^ *SHMT2* is a key enzyme in the glycine biosynthetic pathway and has been implicated in tumor growth and permitting the shift to glycolytic metabolism seen in hypoxic tumor microenvironments.^66^ Interestingly, although *CD9* expression could be elicited in HBEC ALI cultures in response to IL-13 stimulation [GSE185202]^22^ (log_2_FoldChange=1.46, padj=3.34E-45), *SHMT2* was not IL-13 inducible (log_2_FoldChange=-0.055, padj=0.77), highlighting that mechanisms beyond T2 inflammation alone may drive mTORC1-dependent metabolic reprogramming in airway EpCs.

### mTOR signaling regulates cytokine production in CRSwNP basal EpCs

A recent study by Chen et al. reported that glycolysis is required for *ex vivo* production of inflammatory mediators such as CXCL8 by CRS EpCs in response to IL-1β and TNF-α.^53^ Building upon this, we found that the *in vivo* transcriptomic glycolysis score in CRSwNP basal EpCs correlated positively with expression of the chemokines *CXCL8* and *CXCL2* and the alarmin *IL33* (p<0.05) (**Fig 3A**), while there was no significant correlation with *TSLP* (p=0.094) (**Fig 3A**). Having established the close relationship between glycolysis and mTORC1 signaling in CRSwNP EpCs (**Fig 2D**), we hypothesized that mTOR signaling was also necessary for production of these EpC mediators in CRSwNP. To assess this, we expanded primary human BCs from patients with CRSwNP in ex vivo culture, stimulated them with or without the combination of IL-1β and TNF-α and assessed for expression of the indicated cytokines by immunofluorescence. We found that production of CXCL8 and IL-33 was low at baseline, induced by IL-1β and TNF-α, and completely inhibited by the mTORC1/C2 inhibitor Torin 1 (**Fig 3B**, **3C**). CXCL2 expression was present at baseline and did not clearly increase with stimulation, but baseline expression was also inhibited by Torin 1. The Torin 1 inhibitor was not toxic to cells, as stimulated cells robustly upregulated TSLP in the presence or absence of the inhibitor. In summary, here we demonstrate that mTOR signaling regulates the production of several epithelial cytokines predicted to elicit both T2 and non-T2 inflammation, highlighting the potential importance of epithelial mTOR signaling in disease pathogenesis.

**Fig 3:**
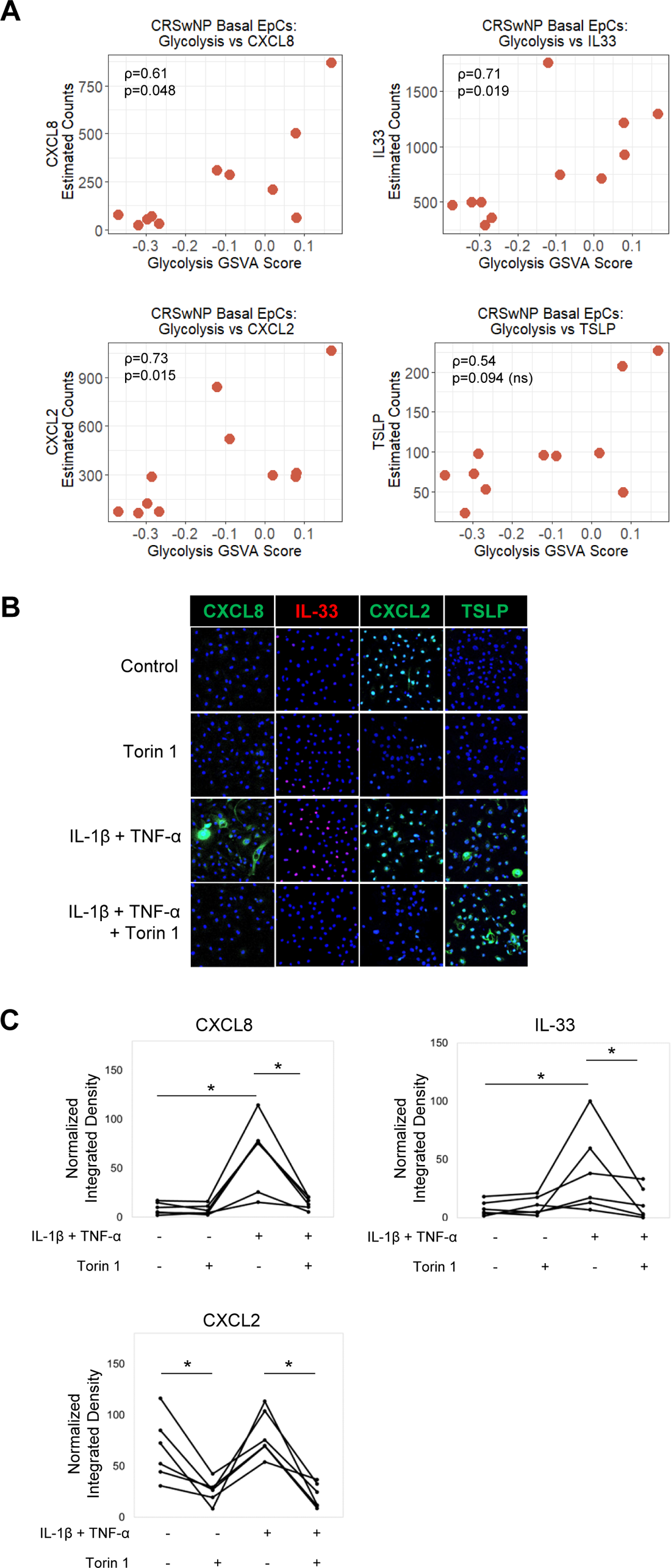
mTOR signaling regulates cytokine production in CRSwNP basal EpCs. A) Scatterplots of glycolysis GSVA score vs estimated counts of selected epithelial cytokines in CRSwNP basal EpCs. *ρ* indicates Spearman’s *rho*. B) Representative images of cytokine and DAPI staining immunocytochemistry in CRSwNP basal EpCs in the presence vs absence of stimulation (IL-1β and TNF-α) and Torin 1 (mTOR inhibitor). Blue represents DAPI staining. C) Quantification of cytokine immunofluorescence for CRSwNP basal EpCs from B. Normalized integrated density refers to the cytokine integrated density divided by the number of nuclei per high power field. The non-parametric Wilcoxon test was used for statistical testing, with pairing by donor.

### The epithelial mTORC1-glycolysis axis correlates with immune cell tissue infiltration in CRSwNP

As we had determined that the mTOR-glycolysis axis was necessary for production of several inflammatory mediators by CRSwNP basal EpCs (**Fig 3**), we asked whether glycolysis in CRSwNP basal EpCs correlated with the presence of immune cells in nasal polyp tissue. To answer this, we explored the Wang scRNA-seq dataset which includes >70,000 cells (immune cells and EpCs) from healthy controls (n=5), CRSsNP (n=5), non-eosinophilic CRSwNP (neCRSwNP, n=5), and eosinophilic CRSwNP (eCRSwNP, n=6).^14^ This dataset was iteratively subclustered into various epithelial and immune cell lineages (**Fig E4**).

As expected, eCRSwNP basal EpCs demonstrated higher glycolysis (p=0.0087) and mTORC1 signaling (p=0.0043) scores than healthy control basal EpCs (**Fig 4A**). Notably, the glycolysis and mTORC1 signaling scores in neCRSwNP were not significantly different from control, and there was a trend toward higher glycolysis and mTORC1 signaling scores in eCRSwNP compared to neCRSwNP, but this was not statistically significant with the limited sample size (neCRSwNP n=5, eCRSwNP n=6) (**Fig 4A**). As we saw previously, the glycolysis score in CRSwNP basal EpCs correlated tightly with mTORC1 signaling (ρ=0.63, p=0.044) (**Fig 4B**).

Whereas IL-33 is recognized for its capacity to promote a T2 immune response through its actions on T_H_2 cells, ILC2s, mast cells, and eosinophils;^67^ CXCL-8 (IL-8) and CCL2 (MCP1) are chemokines that attract various granulocytes and monocytes.^68,69^ Thus, we looked to assess inflammatory cell recovery across diverse immunocytes. Of note, the Wang scRNA-seq dataset did not capture eosinophils, as these cells often do not survive processing for scRNA-seq.^70^ Within the CRSwNP samples in the Wang dataset, we observed that the glycolysis score in basal EpCs correlated positively with the infiltration of total immune cells out of all recovered cells per sample *in vivo* (ρ=0.69, p=0.023) (**Fig 4C**), as well as with the infiltration of CD4 Th2 cells out of all recovered cells per sample (ρ=0.84, p=0.0026) (**Fig 4C**). We also observed that the basal EpC glycolysis score correlated positively with the frequency of three ALOX15^+^ macrophage populations per sample – including FCER2^+^ monocyte-macrophages (ρ=0.83, p=0.0031), CCL18^+^ resident tissue resident macrophages (ρ=0.85, p=0.0016), and FN1^+^ activated tissue resident macrophages (ρ=0.77, p=0.0060) (**Fig 4D**) – populations which Wang et al. report play a key role in the T2 immune pathogenesis of eCRSwNP.^14^ In contrast, the basal EpC glycolysis score was not significantly positively correlated with other macrophage populations (p>0.05) and there was in fact a negative correlation with the frequency of C1Q^+^ resting tissue resident macrophages (ρ=-0.75, p=0.010) (**Fig 4D**). Interestingly, we also found that basal EpC glycolysis correlated positively with various CD8 effector T-cell populations, including CD8 Teff cells (ρ=0.80, p=0.0052) and IFN-γ^+^ CD8 Tem cells (ρ=0.69, p=0.023) (**Fig E5A-B, Table E15**). Prior studies have shown that CD8 T-cells in nasal polyp tissue correlate with eosinophilic inflammation,^71–73^ and that IFN-y prevents eosinophil apoptosis and promotes eosinophil mediator generation,^74,75^ suggesting a mechanism by which CD8 T-cells may contribute to T2 disease. In summary, our analysis of the Wang scRNA-seq dataset identified that the degree of epithelial metabolic reprograming in CRSwNP correlates tightly with the tissue infiltration of several types of immunocytes, demonstrating that metabolic reprograming of EpCs may support a proinflammatory epithelial niche in CRSwNP tissue.

### Wound healing and tissue remodeling genes are differentially expressed in CRSwNP vs CRSsNP and correlate with EpC glycolysis

After having observed that mTOR signaling was required for expression of several basal EpC cytokines and that basal EpC glycolysis correlates with immune cell infiltration in CRSwNP, we returned to our original bulk RNA-seq dataset to better understand the non-immune implications of the metabolic reprogramming in CRSwNP EpCs. Thus, we performed an unsupervised analysis within each EpC subset to identify genes which had both a higher expression in CRSwNP than CRSsNP (log2FoldChange>0.58 and padj<0.05) and a positive correlation with glycolysis (ρ>0 and p<0.05) (**Fig 5A, 5C, 5E**). Annotation of the basal EpC DEGs that correlated with glycolysis against the GO Biological Processes database identified over-representation of genes involved in wound healing (padj=0.018) and ECM remodeling (padj=0.025) **(Fig 5B);** while genes involved in peptidase inhibition and regulation of vasculature were over-represented in transitional and differentiated EpC, respectively (padj<0.05) (**Fig 5D, 5F**). The epithelial wound healing response in CRSwNP involves coordination of extracellular matrix remodeling, cell migration, and type 2 epithelial-mesenchymal transition^76,77^ – all of which likely impose a tremendous energetic stress on the EpCs. Taken together, we propose that the metabolic reprograming of EpCs in CRSwNP supports the energetic requirements of wound healing and tissue remodeling that are central to nasal polyposis.

**Fig 4:**
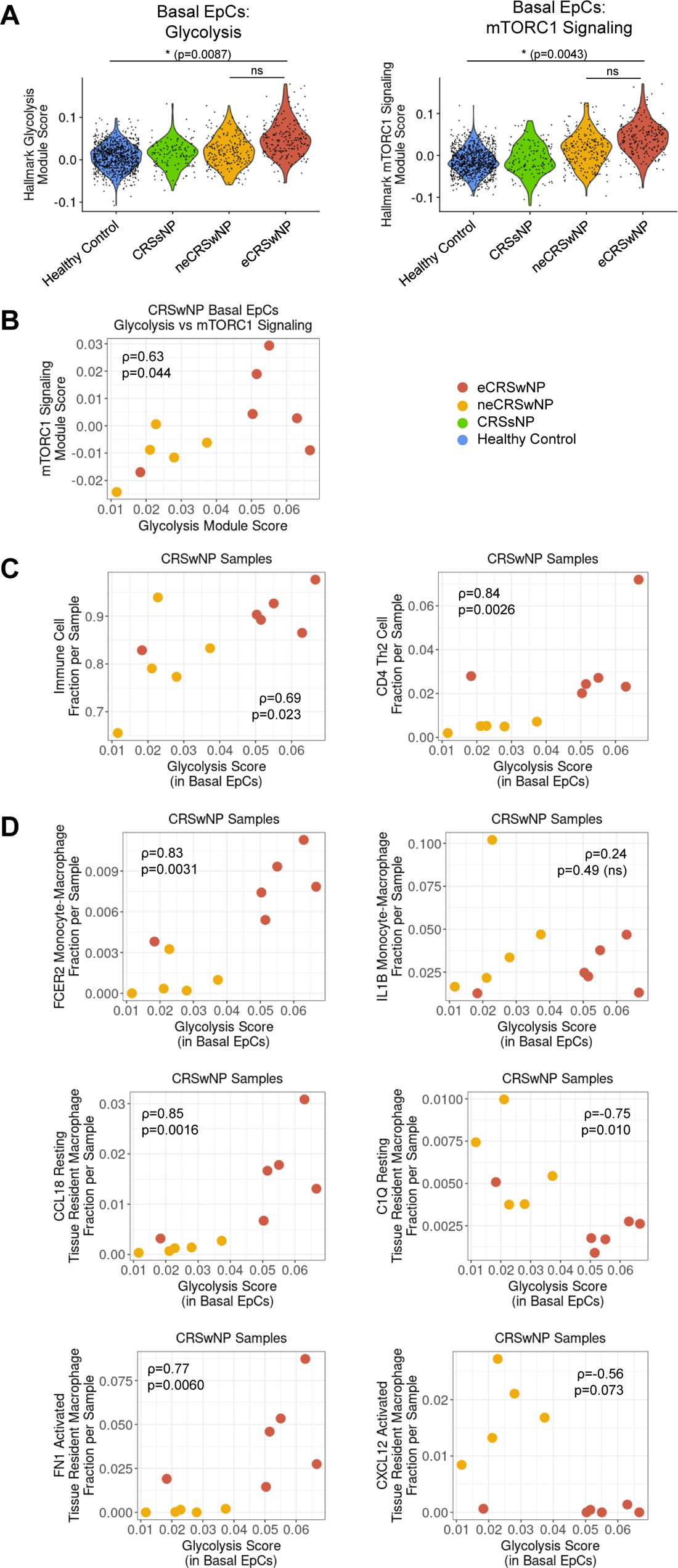
The epithelial mTORC1-glycolysis axis correlates with immune cell tissue infiltration in CRSwNP. A) Violin plots of the glycolysis and mTORC1 signaling scores in basal EpCs from the Wang scRNA-seq dataset. B) Scatterplot of the mean glycolysis score vs mean mTORC1 signaling in basal EpCs from the 11 CRSwNP samples in the Wang scRNA-seq dataset. *ρ* indicates Spearman’s *rho*. C) Scatterplots of the mean glycolysis score in basal EpCs vs fraction of total immune cells and fraction of CD4 Th2 cells recovered by scRNA-seq from the 11 CRSwNP samples in the Wang scRNA-seq dataset. D) Scatterplots of the mean glycolysis score in basal EpCs vs macrophage populations recovered by scRNA-seq from the 11 CRSwNP samples in the Wang scRNA-seq dataset. FCER2^+^ monocyte-macrophages, CCL18^+^ resting tissue resident macrophages, and FN1^+^ resting tissue resident macrophages are each ALOX15^+^ macrophage populations.

**Fig 5:**
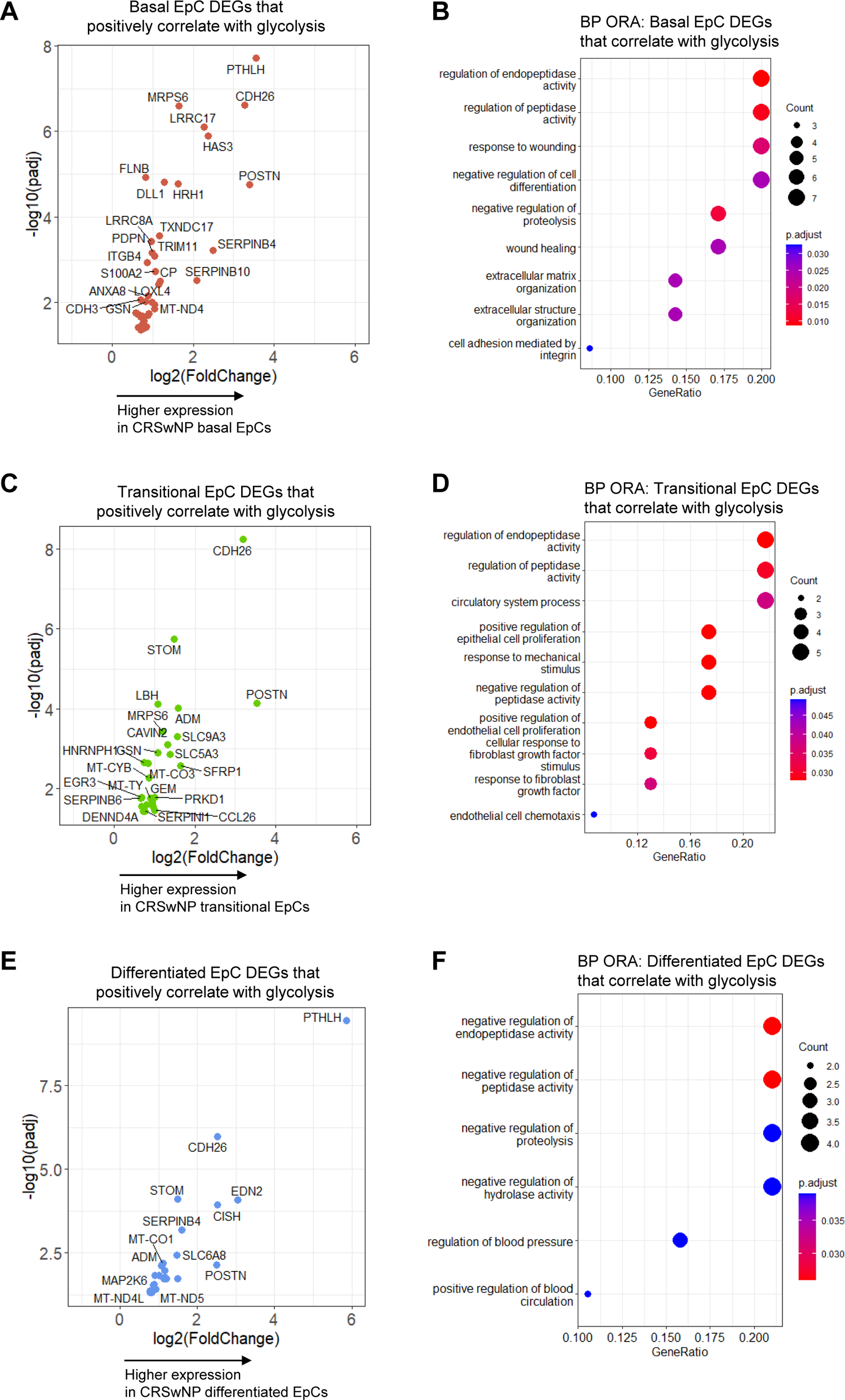
Increased wound healing in CRSwNP correlates with glycolysis. A) Volcano plot of basal EpC DEGs (log_2_FoldChange>0.58 and padj<0.05) with higher expression in CRSwNP that also correlate positively with basal EpC glycolysis (*ρ*>0 and p<0.05). B) Dot plot of top over-represented gene sets (in the gene ontology biological processes database) for basal EpC DEGs with higher expression in CRSwNP that correlate with basal EpC glycolysis. C) Volcano plot of transitional EpC DEGs (log_2_FoldChange>0.58 and padj<0.05) with higher expression in CRSwNP that also correlate positively with transitional EpC glycolysis (*ρ* >0 and p<0.05). D) Dot plot of top over-represented gene sets (in the gene ontology biological processes database) for transitional EpC DEGs with higher expression in CRSwNP that correlate with transitional EpC glycolysis. E) Volcano plot of differentiated EpC DEGs (log_2_FoldChange>0.58 and padj<0.05) with higher expression in CRSwNP that also correlate positively with differentiated EpC glycolysis (ρ>0 and p<0.05). F) Dot plot of top over-represented gene sets (in the gene ontology biological processes database) for differentiated EpC DEGs with higher expression in CRSwNP that correlate with differentiated EpC glycolysis.

### Epithelial glycolysis can be induced by T2 and T17 cytokines *in vitro*, and correlates with T2 cytokine response *in vivo*

Metabolic rewiring in T-cells towards glycolysis can be driven by cytokines and has been long recognized as a central feature of T-cell effector polarization,^78–81^ while more recent studies have begun to explore the drivers and consequences of metabolic reprogramming in airway EpCs.^53,82^ To understand how the local cytokine milieu influences glycolysis in airway EpCs, we analyzed an external bulk RNA-seq dataset of human bronchial EpC (HBEC) air-liquid interface (ALI) cultures which were stimulated with IFN-α, IFN-γ, IL-13, or IL-17 [GSE185202]^22^. Hallmark GSEA of DE testing results for each cytokine vs control demonstrated that the glycolysis gene set was weakly positively enriched in IL-13-stimulated ALIs (padj=0.034) and more strongly positively enriched in IL-17-stimulated ALIs (padj=4.1E-4) (**Fig 6A, Tables E6-7**); in contrast, stimulation with IFN-α and IFN-γ did not lead to significant enrichment of the glycolysis gene set (padj>0.05) (**Fig 6A**). When we examined the cytokine responses of individual HBEC donors using GSVA, stimulation with IL-17 consistently drove an increase in the glycolysis score while the response to IL-13 was variable (**Fig 6B)**.

**Fig 6:**
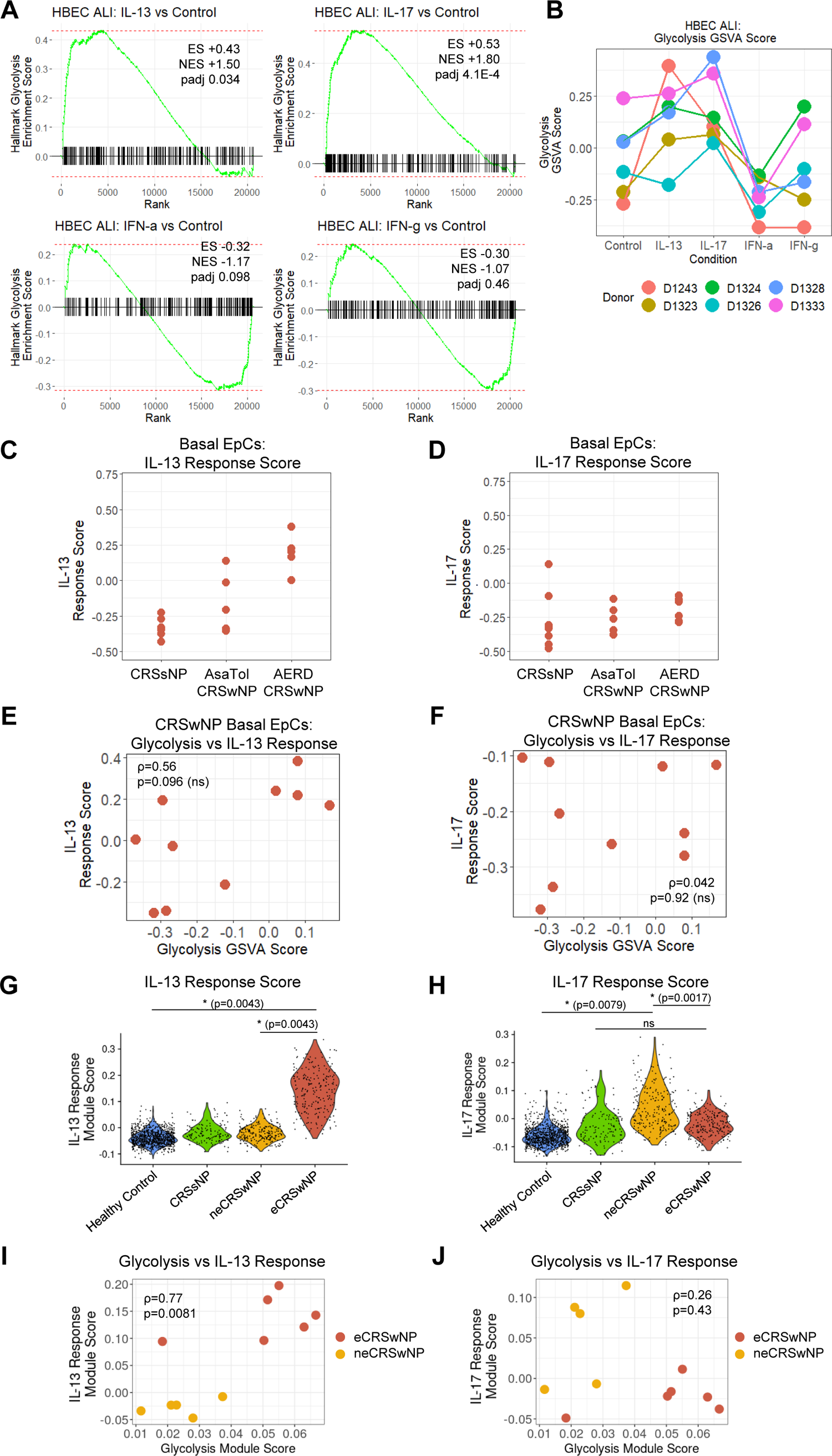
Epithelial glycolysis can be induced by T2 and T17 cytokines. A) GSEA plots of the Hallmark glycolysis gene set for bulk RNA-seq DE testing of HBEC ALI cultures from healthy lung transplant donors stimulated with IL-13, IL-17, IFN-α, or IFN-γ (GSE185202). Positive ES or NES indicates enrichment with cytokine stimulation. B) GSVA glycolysis scores for HBEC ALIs by cytokine stimulation condition. C) IL-13 response score in basal EpCs by disease (including aspirin tolerant CRSwNP and AERD). One sample (from D438) was excluded because the donor had received dupilumab. D) IL-17 response score in basal EpCs by disease (including aspiring tolerant CRSwNP and AERD). E) Scatterplot of glycolysis GSVA score vs IL-13 response score in CRSwNP basal EpCs. ρ indicates Spearman’s *rho*. One sample (from D438) was excluded because the donor had received dupilumab. F) Scatterplot of glycolysis GSVA score vs IL-17 response score in CRSwNP basal EpCs. G) Violin plot of the IL-13 response scores in basal EpCs from the Wang scRNA-seq dataset. Each dot represents one basal EpC. The non-parametric Mann-Whitney U test was performed using the mean score for each donor (control n=5, CRSsNP n=5, neCRSwNP n=5, eCRSwNP n=6). Relevant statistical results are labeled where * denotes p<0.05 and *ns* denotes p>0.05. H) Violin plot of the IL-17 response scores in basal EpCs from the Wang scRNA-seq dataset. I) Scatterplot of the mean glycolysis score vs mean IL-13 response scores in basal EpCs from the 11 CRSwNP samples in the Wang scRNA-seq dataset. ρ indicates Spearman’s *rho*. J) Scatterplot of the mean glycolysis score vs mean IL-17 response scores in basal EpCs from the 11 CRSwNP samples in the Wang scRNA-seq dataset.

We utilized the top 200 genes induced by each cytokine in HBEC ALIs as cytokine response signatures with which we scored basal EpCs in our *in vivo* dataset using GSVA (**Table E10**). As expected, samples from subjects with CRSwNP exhibited higher IL-13 response scores than those with CRSsNP, and within CRSwNP the subjects with AERD exhibited the highest IL-13 response scores (**Fig 6C**). In contrast, the IL-17 response scores were low across CRSsNP and CRSwNP throughout our dataset (**Fig 6D**). While we observed a non-significant trend toward positive correlation between glycolysis and IL-13 response score in CRSwNP basal EpCs (ρ=0.56, p=0.096) (**Fig 6E**), there was no detectable correlation between glycolysis and the low IL-17 response score in CRSwNP EpCs (ρ=0.042, p=0.92) (**Fig 6F**). Here, a caveat is that the lack of association between EpC glycolysis and IL-17 response may be due to the limited diversity in IL-17 response scores in our CRSwNP dataset.

In the Western hemisphere, CRSwNP is predominantly a T2-high and eosinophilic disease,^83,84^ consistent with the elevated transcriptional IL-13 response scores we observed in our bulk RNA-seq dataset (**Fig 6C**). However, CRSwNP in Asia is noted to be more heterogeneous, with neutrophilic and eosinophilic subtypes, as reviewed in **Fig 4**.^83^ Accordingly, in order to examine the relationship between cytokine response and metabolic rewiring across diverse disease subtypes including eCRSwNP and neCRSwNP, we explored this in the Wang scRNA-seq dataset. Within basal EpCs in the Wang dataset, we observed that eCRSwNP samples exhibited the highest IL-13 response score (**Fig 6G**). Similar to our findings in bulk RNA-seq (**Fig 6D**), there was no significant difference in IL-17 response score between CRSsNP and eCRSwNP in the Wang dataset (p>0.05) (**Fig 6H**). In contrast, basal EpCs from neCRSwNP (defined by the authors as fewer than 10 Eos/HPF on nasal polyp histology)^14^ demonstrated modest elevations in the IL-17 response score compared to healthy controls (p=0.0079) (**Fig 6H**), although the IL-17 response score did not significantly correlate with the fraction of neutrophils recovered by scRNA-seq per sample (p=0.066) (**Fig E5C**), consistent with prior reports.^85^ Basal EpCs from neCRSwNP also exhibited modest but statistically significant elevations in IFN-α signaling (p=0.0032) and IFN-γ signaling (p=0.0079) compared to healthy controls (**Fig E6**). Among CRSwNP samples in the Wang dataset, the glycolysis score in CRSwNP basal EpCs correlated tightly with IL-13 response (ρ=0.77, p=0.0081) (**Fig 6I**) but not with the IL-17 response (ρ=0.26, p=0.43) (**Fig 6J**). Here, although not significant within only 5 samples, neCRSwNP samples with higher IL-17 response scores tended to have higher glycolysis scores (**Fig 6J**), suggesting that non-T2 cytokines such as IL-17 could potentially drive metabolic reprogramming in upper airway EpCs. Taken together, we observed that the enhanced glycolytic activity in CRSwNP EpCs correlates with the T2 cytokine response *in vivo*.

Finally, to investigate the relationship between T2 inflammation and EpC glycolysis more closely, we queried the pan-epithelial DEGs with higher expression in CRSwNP (**Fig 2H**) against the list of genes induced by IL-13 stimulation to identify which pan-epithelial DEGs could be attributable to IL-13 response and which could not. We found that 209 of the 359 pan-epithelial DEGs with higher expression in CRSwNP (over 50%) were not directly IL-13 responsive (**Fig E7A**). Interestingly, whereas *PFKP* was IL-13 inducible (log_2_FoldChange=1.18, padj=5.02E-23) in the HBEC ALIs, *SLC2A1* was not (log_2_FoldChange=-0.14, padj=0.33). This indicates that metabolic reprograming in CRSwNP EpCs cannot be fully attributed to direct effects of IL-13 signaling, and that additional mechanisms are also present. Interestingly, over-representation analysis of the 209 non-IL13-responsive pan-epithelial DEGs identified gene sets involved in type 1 interferon signaling (padj=0.043) (**Fig E7B**), highlighting a contribution of non-T2 mechanisms in the immune pathogenesis of CRSwNP.

### Glycolysis and mTORC1 signaling are enriched in CRSwNP across basal and secretory EpCs as compared to healthy controls

As our flow cytometric strategy did not distinguish between secretory and ciliated EpCs, we next assessed the mTORC1-glycolysis axis in a recently reported scRNA-seq dataset that included over >100,000 ethmoid sinus EpCs from donors with CRSwNP (n=5) and healthy controls (n=4).^13^ As expected, gene module scoring demonstrated higher expression of the IL13 response score (p=0.016) (**Fig 7A**), Hallmark glycolysis gene set (p=0.016) (**Fig 7B**), and Hallmark mTORC1 signaling gene set (p=0.016) (**Fig 7C**) in basal EpCs from CRSwNP vs healthy controls. Although limited to 5 samples with CRSwNP, there was still a trend toward positive correlation between glycolysis and mTORC1 signaling in CRSwNP basal EpCs (ρ=0.70, p=0.23) (**Fig 7D**). Increased expression of key glycolytic genes including *SLC2A1* was evident in the CRSwNP basal EpCs (**Fig 7E**).

**Fig 7:**
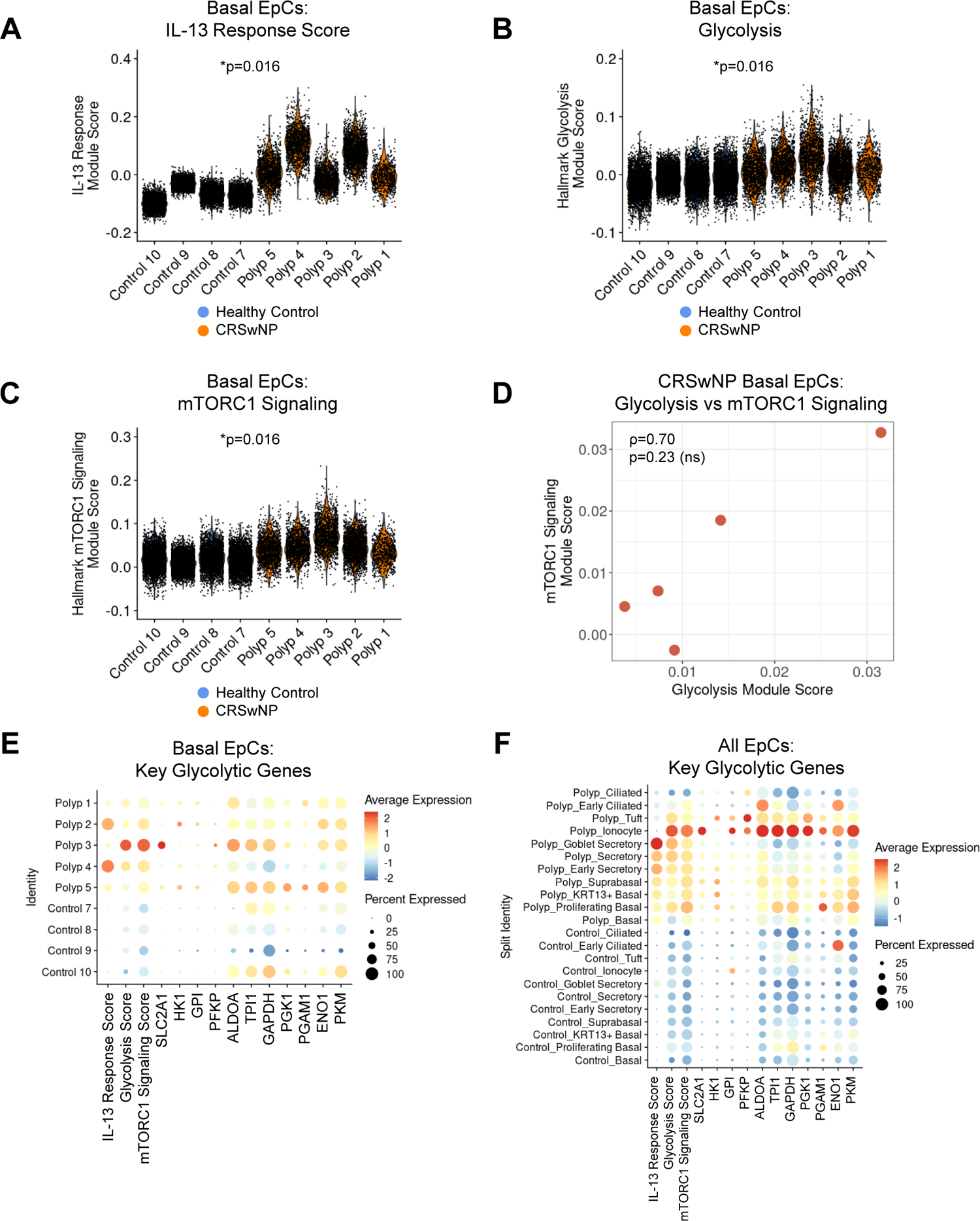
Glycolysis and mTOR signaling are enriched in CRSwNP across basal and secretory EpCs. A) Violin plot of the IL-13 response score in basal EpCs from the Kotas scRNA-seq dataset. Each dot represents one basal EpC. The non-parametric Mann-Whitney U test was performed using the mean score for each donor (control n=4, CRSwNP n=5). B) Violin plot of the glycolysis score in basal EpCs from the Kotas scRNA-seq dataset. C) Violin plot of the mTORC1 signaling score in basal EpCs from the Kotas sRNA-seq dataset. D) Scatterplot of the mean glycolysis score vs mean mTORC1 signaling score in basal EpCs from the 5 CRSwNP samples in the Kotas scRNA-seq dataset. ρ indicates Spearman’s *rho*. E) Dot plot of module scores and key genes in the glycolysis pathway for basal EpCs in the Kotas scRNA-seq dataset, split by donor. F) Dot plot of module scores and key genes in the glycolysis pathway for various EpC lineages in the Kotas scRNA-seq dataset, split by disease (control and CRSwNP).

The mTORC1-glycolytic axis was apparent in CRSwNP vs control across multiple lineages of EpCs, including basal, suprabasal, secretory, and goblet secretory EpCs (**Fig 7F**); by contrast, this wasn’t appreciable in ciliated EpCs. Interestingly, enrichment of mTORC1 signaling and glycolytic genes were highly apparent in CRSwNP ionocytes compared to healthy control ionocytes (**Fig 7F**), although the CRSwNP ionocytes did not exhibit strong IL-13 response scores. The low ionocyte IL-13 response scores may reflect a lack of ionocytes in the ALI cultures used to define the IL-13 response signature or reflect a role for non-T2 mechanisms in regulating mTORC1-associated metabolic reprogramming in CRSwNP EpCs. Similarly, CRSwNP goblet secretory EpCs with high IL-13 response score exhibited little upregulation of glycolytic genes (**Fig 7F**), demonstrating that T2 cytokine signaling alone is not sufficient to elicit the full cascade of glycolytic rewiring that was detected in basal and suprabasal EpCs.

Having established that metabolic rewiring of EpCs may support tissue remodeling in CRSwNP (**Fig 5**), we queried if this axis was related to compositional changes among EpCs in the Kotas dataset. Although limited to only 5 CRSwNP samples, we observed the basal EpC glycolysis score was negatively correlated with the fraction of KRT13+ “hillock” basal EpCs out of all cells recovered per sample (ρ=1, p=0.017) (**Fig E8**). The function of hillock basal EpCs is poorly understand and the authors of the Kotas dataset did not identify any differences in their frequency between CRSwNP and healthy control,^13^ but it has been proposed that they have roles in squamous metaplasia and immunomodulatory functions as well.^86–88^

### The mTORC1-glycolysis axis is detected in the lower airway and correlates with T2 and non-T2 cytokine response genes

Finally, to understand whether the inflammatory mTOR-glycolysis axis is also present in the lower airway, we analyzed the Immune Mechanisms of Severe Asthma (IMSA) bulk RNA-seq dataset of bronchial brushings (containing both EpCs and immune cells) from 65 adults (17 healthy controls, 25 with mild-to-moderate asthma, and 23 with severe asthma).^39,40^ DE analysis of severe vs mild-to-moderate asthma (while controlling for batch, sex, and age as covariates) identified 96 DEGs (|log_2_FoldChange|>0.58 and padj<0.05), including 86 DEGs with higher expression in bronchial brushings from severe asthma (**Fig 8A**).

**Fig 8:**
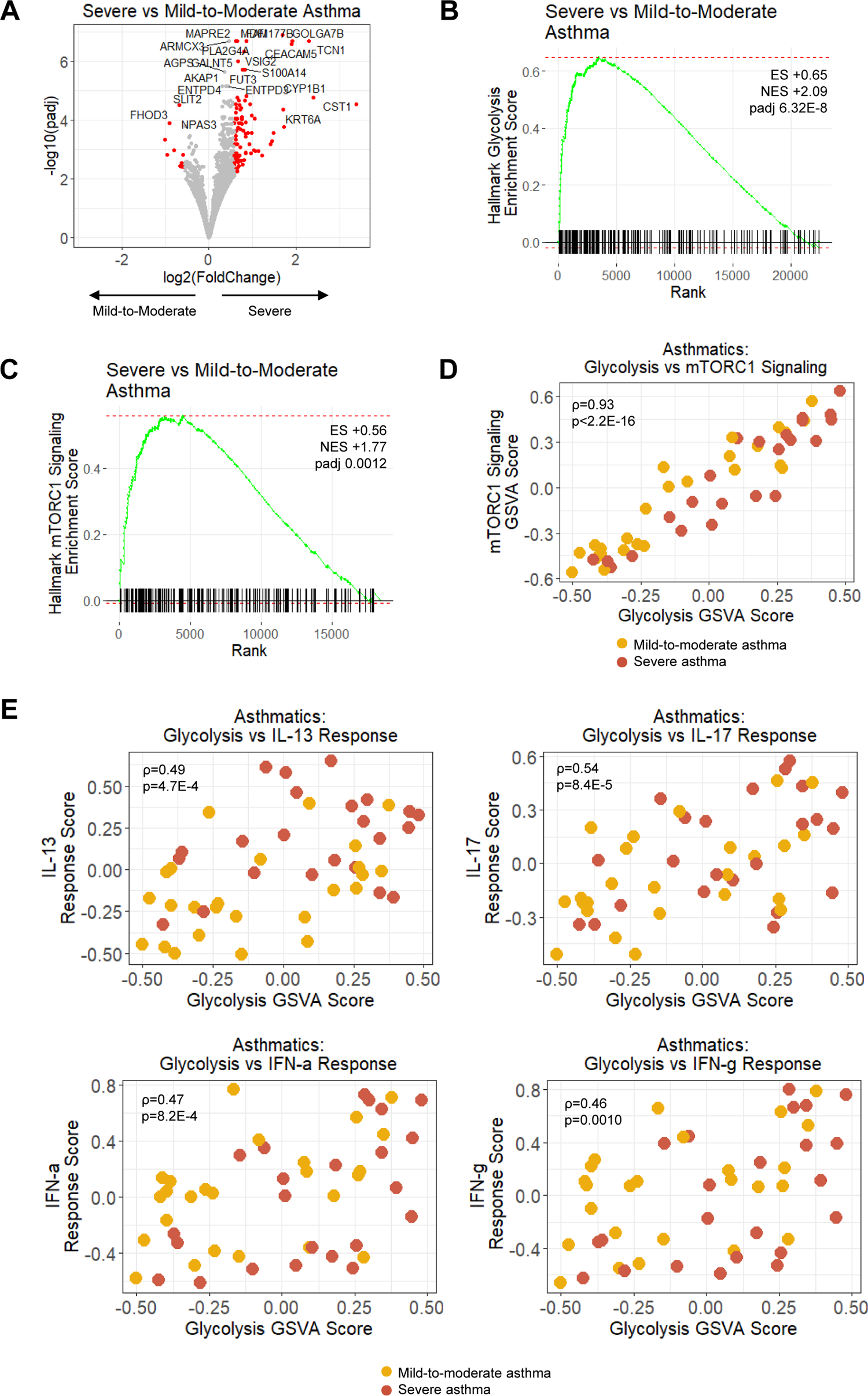
The mTOR-glycolysis axis correlates with T2 and non-T2 cytokine response genes in asthma. A) Volcano plot of severe asthma vs mild-to-moderate asthma DE testing in the IMSA bronchial brushing bulk RNA-seq dataset [GSE158752]. Positive log_2_FoldChange indicates higher expression in severe asthma. Red indicates DEGs meeting |log_2_FoldChange|>0.58 and padj<0.05. B) GSEA plot of the Hallmark glycolysis gene set for bulk RNA-seq DE testing of severe vs mild-to-moderate asthma. Positive ES or NES indicates enrichment in severe asthma. C) GSEA plot of the Hallmark mTORC1 signaling gene set for bulk RNA-seq DE testing of severe vs mild-to-moderate asthma. D) Scatterplot of the glycolysis score vs mTORC1 signaling score in asthmatic bronchial brushings in the IMSA bulk RNA-seq dataset. ρ indicates Spearman’s *rho*. E) Scatterplots of the glycolysis score vs IL-13, IL-17, IFN-α, and IFN-γ response scores in asthmatic bronchial brushings in the IMSA bulk RNA-seq dataset.

GSEA of the Hallmark gene sets identified positive enrichment of both glycolysis (padj=6.32E-8) (**Fig 8B**) and mTORC1 signaling (padj=0.0012) (**Fig 8C**) in severe asthma. Furthermore, GSVA scoring for these gene sets revealed that glycolysis and mTORC1 signaling were tightly positively correlated in asthmatic bronchial brushings (ρ=0.93, p<2.2E-16) (**Fig 8D**). Asthma, particularly severe asthma, represents a heterogeneous disease with evidence of combinatorial variants of T1, T2, and T17 inflammation;^39,89–91^ thus we wondered if the local cytokine milieu was also related to epithelial metabolic reprogramming in asthmatic tissue. We found that the glycolysis score was weakly but positively correlated with multiple cytokine response signatures in asthmatic bronchial brushings *in vivo* (IL-17 ρ=0.54, p=8.4E-5; IL-13 ρ=0.49, p=4.7E-4) (**Fig 8E**), consistent with our findings in HBEC ALI cultures (**Fig 6A**). Thus, in the lower airway *in vivo,* epithelial metabolic reprograming may reflect a response to stimulation from one (or more) of several cytokines.

## DISCUSSION AND CONCLUSIONS

Defining the pathways that maintain barrier tissue inflammation is essential to understanding chronic respiratory diseases such as CRSwNP. A recent seminal paper by Chen and colleagues used metabolomics from nasal secretions and measurements of extracellular acidification in stimulated epithelial cells to identify enhanced glycolysis in nasal epithelial cells from CRS.^53^ Additionally, they demonstrated that glycolysis is required for the production of several EpC cytokines, including IL-1α, TNF-α, IL-1β, CXCL8, and CCL20.^53^ In this study, we demonstrate that enhanced glycolytic programming detected in the epithelium of CRSwNP is tightly linked to mTORC1 pathway, and further demonstrate that mTOR regulates airway epithelial cytokine generation. Moreover, we find a close correlation *in vivo* between EpC glycolytic reprogramming and EpC cytokine generation, inflammation, and epithelial remodeling. Taken together, our findings highlight a critical role for mTORC1-dependent metabolic reprogramming of airway EpCs in chronic airway inflammation.

Whereas to date most studies of immunometabolism in chronic respiratory diseases have focused on immune cell populations,^92,93^ these studies highlight the field of epithelial immunometabolism, which is an emerging area of investigation.^94^

Increased mTORC signaling has previously been implicated in chronic airway inflammation. In the upper airway, increased phospho-mTOR staining has been observed in CRSwNP lysates^95,96^. In the lower airway, multiple groups have detected increased phosphorylation of mTORC1 targets in whole lung lysates in murine models of asthma.^97–99^ Accordingly, studies using mTOR inhibitors and mTOR knockout models have begun to explore the functional consequences of mTOR signaling in lung inflammation. Several studies have shown that systemic administration of rapamycin (which inhibits mTORC1 acutely and downregulates mTORC2 chronically^100^) attenuates airway inflammation and airway hyperresponsiveness (AHR) in mouse models of asthma,^97,101,102^ and a recent study found that rapamycin lowered serum levels of IL-4 and IL-17 while restoring IFN-γ in a mouse asthma model.^99^ Similarly, administration of Torin 2 (which inhibits both mTORC1 and mTORC2) led to decreased goblet cell hyperplasia and decreased AHR in a mouse asthma model.^103^However, very few studies have pursued epithelial-specific knockout models to identify the consequences of epithelial mTORC signaling *in vivo*. Interestingly, one study demonstrated that abrogating mTORC1 and mTORC2 signaling in the bronchial epithelium via EpC-specific *mTOR* deletion did not ameliorate airway inflammation in asthmatic mice.^104^ By contrast, a study in the small intestine demonstrated that intestinal EpC-specific deletion of *RAPTOR* (which is required for mTORC1 but not mTORC2 signaling) resulted in impaired formation of tuft cells and decreased T2 immunity in the context of parasitic infection.^105^ Thus, while we do find a correlation between the recovery of Th2 cells and the EpC glycolysis score, more studies are needed to understand the contribution of epithelial mTORC1 and mTORC2 to inflammation *in vivo*.

Beyond perpetuating chronic inflammation, our findings suggest a potential role for epithelial mTORC1 activity in supporting the wound healing response in CRSwNP (**Fig 5**). Previous reports have demonstrated that mTORC1 signaling in murine epidermal EpCs is necessary for and can augment cutaneous wound healing.^106^ Similarly, in the gastrointestinal tract, mTORC1 signaling in murine intestinal enterocytes is critical for epithelial regeneration following surgical or radiation-induced injury and this process can be highjacked in neoplastic tissue.^107–109^ In the human airways, prior reports have identified abnormal wound healing of airway EpCs in chronic respiratory diseases, including increased markers of epithelial-mesenchymal transition (EMT) in CRSwNP tissue^110,111^ and dysfunctional behavior of asthmatic airway EpCs in wound healing assays.^112,113^ Intriguingly, this latter finding suggests that enhanced mTORC1-mediated upregulation of wound healing genes may even be a compensatory mechanism for other defects in tissue repair.

We found that, across stages of secretory EpC development, mTORC1 signaling was tightly correlated with GSVA glycolytic score in bulk RNA-seq and was tightly correlated with glycolytic module score in two additional single cell datasets. One plausible mechanism by which mTORC1 signaling may enhance glycolysis in airway EpCs is through hypoxia-inducible factor (HIF) signaling. HIF signaling is both elicited by mTOR through direct transcriptional and translational mechanisms,^61,62^ and widely recognized to promote expression of glycolytic genes in several physiologic contexts.^114^ Indeed, hypoxia genes were strongly enriched in CRSwNP vs CRSsNP EpCs (**Fig E2**). Furthermore, we noted that *EGLN3,* encoding a key alpha ketoglutarate-dependent hydroxylase that is expressed in response to HIF-1α,^56,115^ was among the top DEGs with higher expression in CRSwNP in basal EpCs (log_2_FoldChange=1.71, padj=9.68E-7) (**Fig 1F**) and across all EpC subsets (log_2_FoldChange=2.94, padj=6.71E-27) (**Fig 1H**). Importantly, an epithelial axis of mTOR signaling driving HIF-1α nuclear localization, enhanced glycolytic metabolism, and glycolytic-dependent wound healing was recently described in the context of IL-17 skin inflammation.^116^ Taken together, these findings suggest that mTOR-dependent glycolytic reprogramming may be a conserved axis in injured epithelium.

Although we observed that EpC glycolysis correlated with an IL-13 response score in CRSwNP (**Fig 6**) *in vivo*, this association does not exclude the presence of concomitant cytokines driving these metabolic alterations in diseased tissue. In HBEC ALI cultures *in vitro*, IL-13 stimulation only weakly drove glycolysis (**Fig 6**) and key glycolytic genes such as *SLC2A1* were not IL-13 inducible (**Fig E7**); therefore T2 cytokines alone may not be sufficient to drive mTORC1-dependent glycolytic reprograming in airway EpCs. Additionally, one study participant (D438) with the AERD variant of CRSwNP had received dupilumab (which antagonizes IL-4 and IL-13 signaling through the alpha chain of the IL-4 receptor) before undergoing sinus surgery. Basal EpCs from subject D438 exhibited low IL-13 response score, as expected, but still demonstrated relatively high glycolysis score and mTORC1 signaling score [**Fig E9**], suggesting that a high IL-13 response score is not required, and that mechanisms other than IL-4/13 signaling can cause metabolic reprograming in CRSwNP EpCs. Moreover, the findings from cultured EpCs that IL-17 can upregulate the glycolysis score (**Fig 6A-B**), and that mTOR-dependent EpC cytokines can be elicited by TNF-α and IL-1β (**Fig 3B**), by LPS ^117^, and by flagellin^118^ demonstrate the potential for both endogenous and exogenous insults to elicit changes in EpC metabolic pathways with profound consequences for tissue inflammation and remodeling.

Our *in vivo* data demonstrated a strong correlation between glycolysis and IL-17 response score in the lower airway, but little correlation in the upper airway in eCRSwNP, neCRSwNP, or CRSsNP. Although it is possible that the upper and lower airway epithelium respond differentially to these environmental cues, an equally plausible explanation is that the limited diversity of IL-17 response scores in our CRS datasets precluded our ability to detect non-T2 cues that contribute to mTORC1-dependent metabolic reprogramming in CRSwNP. Furthermore, although we did not observe that EpC glycolysis correlated with an IL-17 response score in CRSwNP *in vivo* (**Fig 6**), we found that IL-17 stimulation more robustly drove glycolytic metabolism in HBEC ALIs *in vitro* (**Fig 8**), consistent with the mTOR-dependent induction of glycolysis seen by others in cutaneous EpCs.^116^

Finally, a limitation of this study is that we used epithelial transcriptomics to assess the glycolytic behavior of CRS EpCs. Although studies in the field of cancer biology have embraced the use of transcriptomic glycolysis scores for clinical prognostication in epithelial tumors,^119–121^ further studies are needed to understand how closely transcriptomic readouts mirror metabolism in chronic inflammation. While our analysis lacks the resolution of metabolomics and we were limited to assessing transcriptional signatures rather than individual glycolytic intermediates, our RNA-seq findings are highly consistent with those reported by Chen et al. who identified enhanced glycolysis in CRSwNP EpCs through assessing bioenergetic function with the Seahorse assay and performing metabolomic analyses.^53^ In addition, we replicated the finding of an increased glycolytic transcriptional signature in CRSwNP EpCs in 2 independent scRNA-seq cohorts.

Here we have shown that mTORC1 activity in the epithelium is upregulated in CRSwNP, as compared to CRSsNP, correlates with epithelial glycolytic reprogramming, and regulates airway EpC cytokine generation. Furthermore, we find that epithelial glycolytic pathways are closely correlated with both T2 and non-T2 immune cell recovery, and with the epithelial expression of wound healing genes. As mTORC1 signaling is a master regulator of cell growth and cell fate, pairing alterations in nutrient metabolism with protein synthesis required for tissue development and repair, these findings suggest that mTORC1 may play a key role in the maintenance of barrier integrity, repair, and remodeling in CRSwNP, and identify mTORC1-dependent pathways as targets for further study.

## Supporting information

Online Repository

## Abbreviations

T2: type 2

T2I: inflammation

CRSwNP: chronic rhinosinusitis with nasal polyps

CRSsNP: chronic rhinosinusitis without nasal polyps

AERD: aspirin-exacerbated respiratory disease

EpCs: epithelial cells

IL-4: interleukin-4

IL-13: interleukin-13

RNA-seq: RNA-sequencing

scRNA-seq: single cell RNA-sequencing

DE: differential expression

DEGs: differentially expressed genes

BCAM: basal cell adhesion molecule

KRT5: keratin 5

TP63: tumor protein 63

NGFR: nerve growth factor receptor

EpCAM: epithelial cell adhesion molecule

SCGB1A1: secretoglobin family 1A

MUC5AC: mucin 5AC

IRS: insulin receptor substrate

mTOR: mammalian target of rapamycin

mTORC: mTOR complex

ILC: innate lymphoid cell

IRS: insulin receptor substrate

HIF-1α: hypoxia-inducible factor 1α

EMT: epithelial-mesenchymal transition

## AUTHOR CONTRIBUTIONS

Conceptualization: GXH, NRH, MGA, NAB

Collection of human samples: AZM, RER, RWB, NB, JH, TR, KMB, TML

Human single cell and bulk sequencing analysis: GXH, MVM, NRH, MGA, NAB

Human *ex vivo* experiments and imaging: KZ, ML

Funding acquisition: SEW, AR, JEG, TSH, JAB, NAB

Project administration: NAB

Supervision: MGA, NAB

Writing – original draft: GXH, MAG, NAB

Writing – review & editing: GXH, MAG, NAB, JEG

