## Supplementary material for "Increased epithelial mTORC1 activity in chronic rhinosinusitis with nasal polyps": Online Repository

1. Supplementary Figures and Legends
    a. Fig E1: Overlap in top CRSwNP vs CRSsNP DEGs in transitional and differentiated EpCs
    b. Fig E2: Hallmark GSEA results for pan-epithelial CRSwNP vs CRSsNP DE testing
    c. Fig E3: Glycolysis was not significantly correlated with mTORC2 transcriptional activity in CRSwNP EpCs
    d. Fig E4: Clustering and sub-clustering of the Wang scRNA-seq data set
    e. Fig E5: Other correlations of EpC gene expression scores with immune cell tissue infiltration in the Wang dataset
    f. Fig E6: Interferon response scores are elevated in neCRSwNP basal EpCs
    g. Fig E7: Genes involved in interferon signaling are over-represented among non-IL-13-responsive pan-epithelial DEGs
    h. Fig E8: EpC glycolysis negatively correlates with the fraction of KRT13+ “hillock” basal EpCs in CRSwNP
    i. Fig E9: Glycolysis and mTORC1 signaling were preserved despite dupilumab administration in one subject with AERD (D438)
2. Excel file with data Tables E1-E20.
    a. Table E1: Metadata for bulk RNA-seq study subjects
    b. Table E2: Top DEGs for CRSwNP vs CRSsNP in basal EpCs
    c. Table E3: Top DEGs for CRSwNP vs CRSsNP in transitional EpCs
    d. Table E4: Top DEGs for CRSwNP vs CRSsNP in differentiated EpCs
    e. Table E5: Top pan-epithelial DEGs for CRSwNP vs CRSsNP
    f. Table E6: Top DEGs for IL-13 stimulation vs Control in HBEC ALIs
    g. Table E7: Top DEGs for IL-17 stimulation vs Control in HBEC ALIs
    h. Table E8: Top DEGs for IFN-α stimulation vs Control in HBEC ALIs
    i. Table E9: Top DEGs for IFN-γ stimulation vs Control in HBEC ALIs
    j. Table E10: Genes in epithelial cytokine response signatures
    k. Table E11: Top cluster markers in Wang scRNA-seq dataset for all cells

l. Table E12: Top cluster markers in Wang scRNA-seq dataset for epithelial cells

m. Table E13: Top cluster markers in Wang scRNA-seq dataset for T cells, NK cells, and ILCs

n. Table E14: Top cluster markers in Wang scRNA-seq dataset for CD4 Th2 and ILC2s

o. Table E15: Top cluster markers in Wang scRNA-seq dataset for CD8 Tem and CD8 Tnaive

p. Table E16: Top cluster markers in Wang scRNA-seq dataset for CD8 Teff and NK

q. Table E17: Top cluster markers in Wang scRNA-seq dataset for Tγδ and NK

r. Table E18: Top cluster markers in Wang scRNA-seq dataset for mononuclear phagocytes

s. Table E19: Top cluster markers in Wang scRNA-seq dataset for resting tissue resident macrophages

t. Table E20: Top cluster markers in Wang scRNA-seq dataset for activated tissue resident macrophages

**SUPPLEMENTARY FIGURES**

#### Fig E1

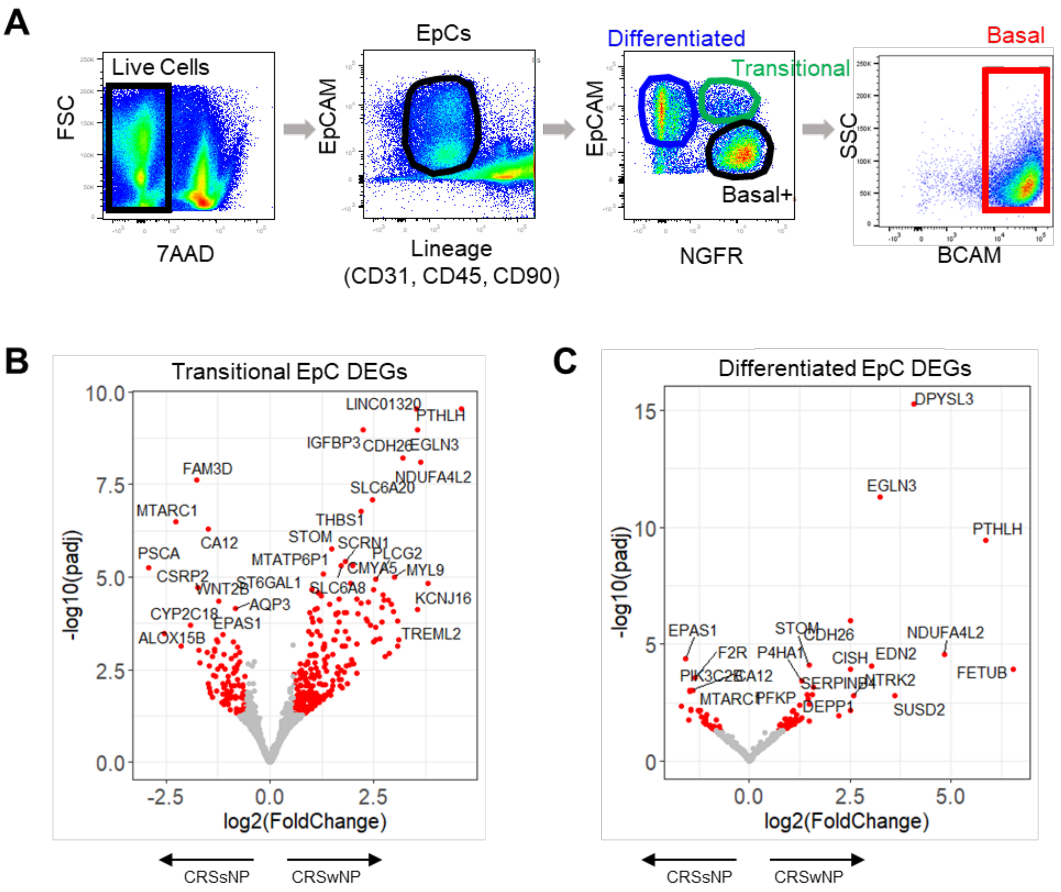


**Fig E1: Overlap in top CRSwNP vs CRSsNP DEGs in transitional and differentiated EpCs**

A) Flow cytometric sorting algorithm for isolation of basal, transitional, and differentiated EpCs

B) Volcano plot of CRSwNP vs CRSsNP DE testing in transitional EpCs. Positive $log_2$FoldChange indicates higher expression in CRSwNP. Red indicates DEGs meeting $|log_2FoldChange|>0.58$ and $padj<0.05$.

C) Volcano plot of CRSwNP vs CRSsNP DE testing in differentiated EpCs.

**Fig E2**

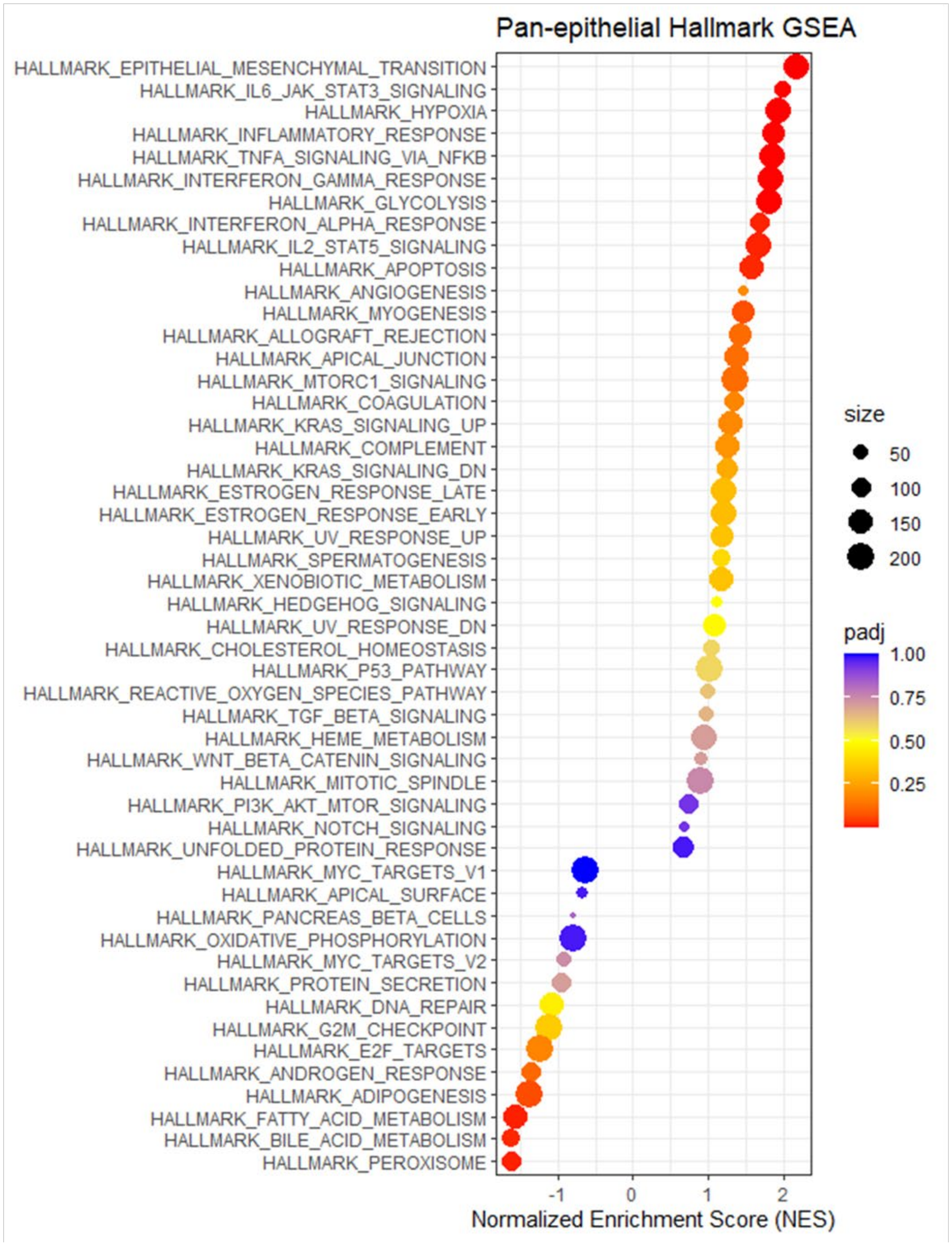


**Fig E2: Hallmark GSEA results for pan-epithelial CRSwNP vs CRSsNP DE testing**

Dot plot of Hallmark GSEA results for CRSwNP vs CRSsNP pan-epithelial DE testing. Positive ES or NES indicates enrichment in CRSwNP.

**Fig E3**

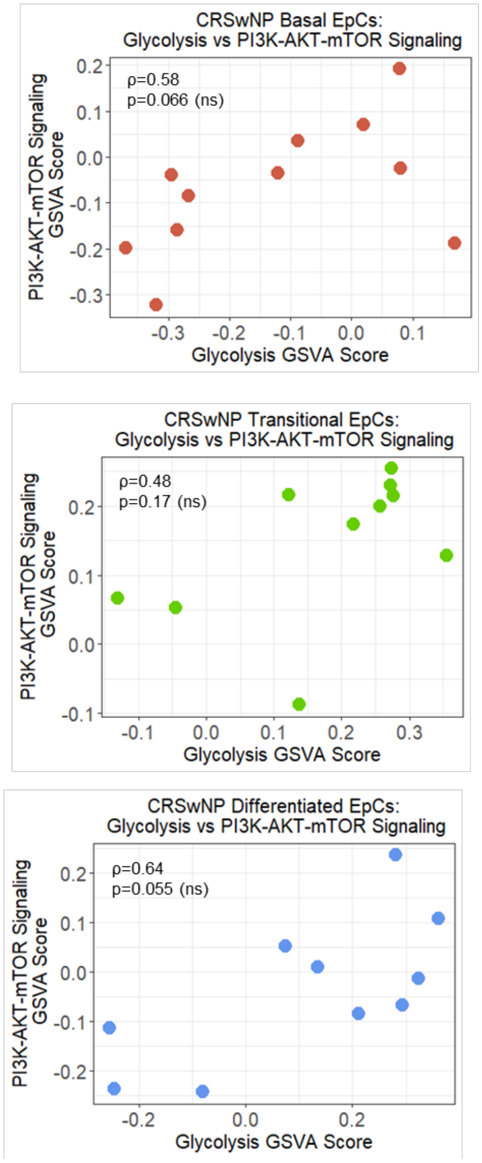


**Fig E3: Glycolysis was not significantly correlated with mTORC2 transcriptional activity in CRSwNP EpCs**

Scatterplots of glycolysis GSVA score vs PI3K-AKT-mTOR signaling GSVA score in CRSwNP basal EpCs, CRSwNP transitional EpCs, and CRSwNP differentiated EpCs. ρ indicates Spearman's *rho*.

#### Fig E4

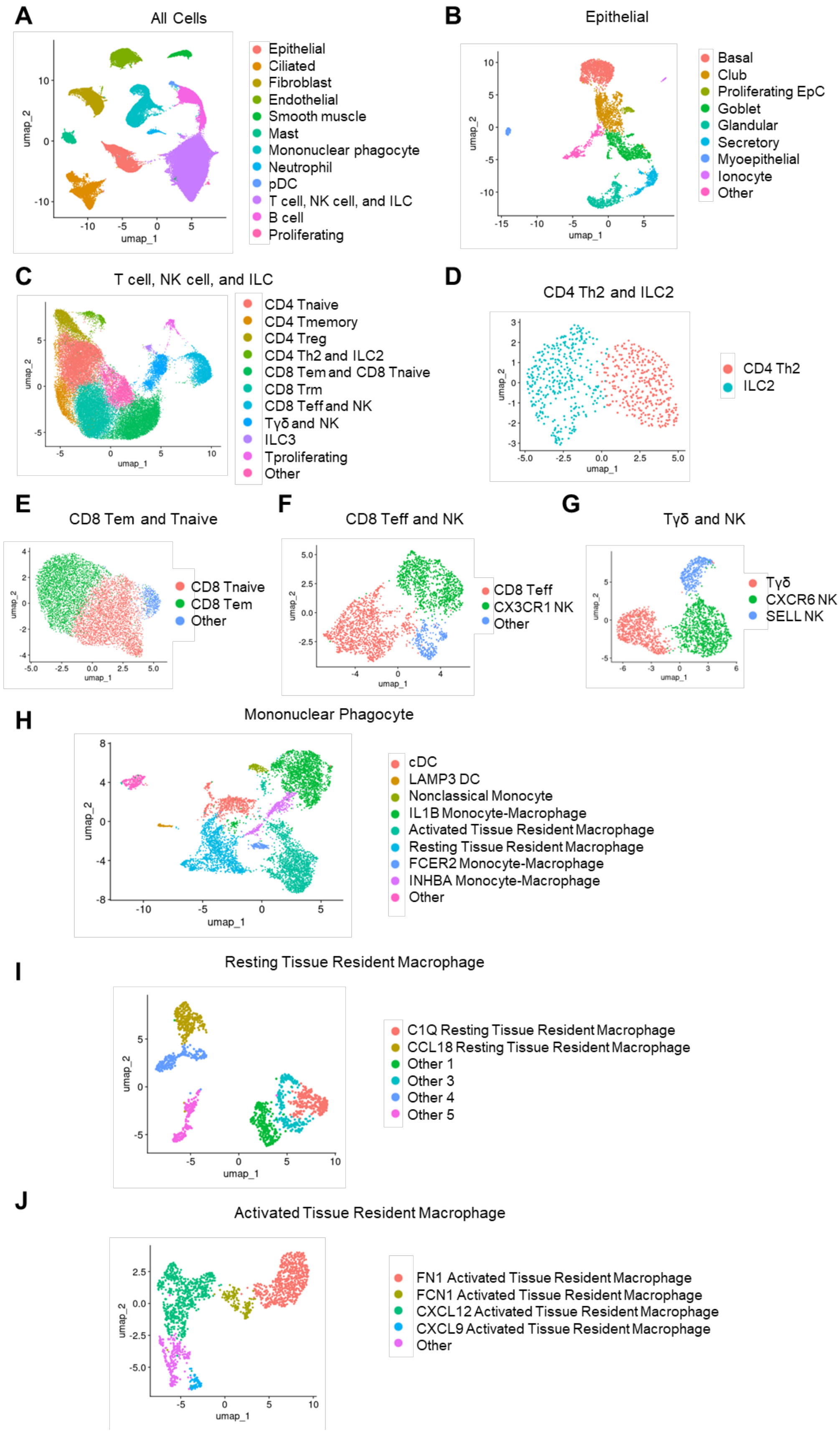


**Fig E4: Clustering and sub-clustering of the Wang scRNA-seq data set**

A) Harmonized UMAP of immune, stromal, and epithelial cells.

B) Subclustered UMAP of EpCs.

C) Subclustered UMAP of T cells, NK cells, and ILCs.

D) Subclustered UMAP of CD4 Th2 cells and ILC2s.

E) Subclustered UMAP of CD8 Tem and CD8 Tnaive cells.

F) Subclusters UMAP of CD8 Teff and NK cells.

G) Subclustered UMAP of Tγδ cells and NK cells.

H) Subclustered UMAP of mononuclear phagocytes (cDCs, monocytes, monocyte-macrophages, and tissue resident macrophages).

I) Subclustered UMAP of resting tissue resident macrophages.

J) Subclustered UMAP of activated tissue resident macrophages.

**Fig E5**

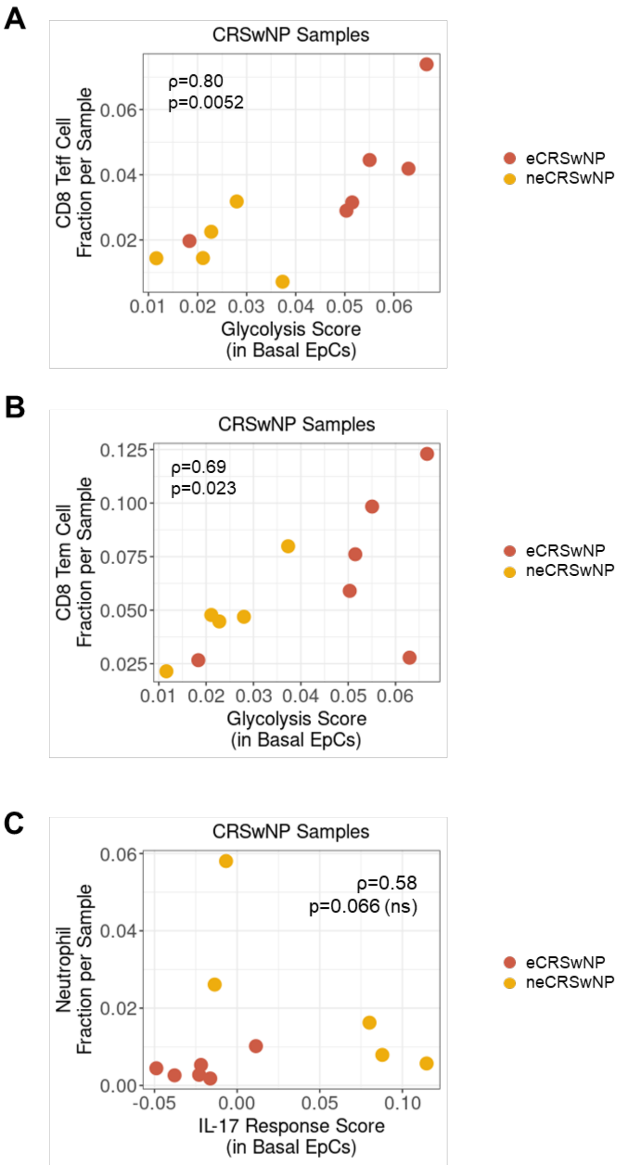

**Fig E5: Other correlations of EpC gene expression scores with immune cell tissue infiltration in the Wang dataset**

A) Scatterplot of the mean glycolysis score in basal EpCs vs fraction of CD8 Teff cells recovered by scRNA-seq from the 11 CRSwNP samples in the Wang scRNA-seq dataset. $\rho$ indicates Spearman's *rho*.

B) Scatterplot of the mean glycolysis score in basal EpCs vs fraction of CD8 Tem cells recovered by scRNA-seq from the 11 CRSwNP samples in the Wang scRNA-seq dataset.

C) Scatterplot of the mean IL-17 response score in basal EpCs vs fraction of neutrophils recovered by scRNA-seq from the 11 CRSwNP samples in the Wang scRNA-seq dataset.

### Fig E6

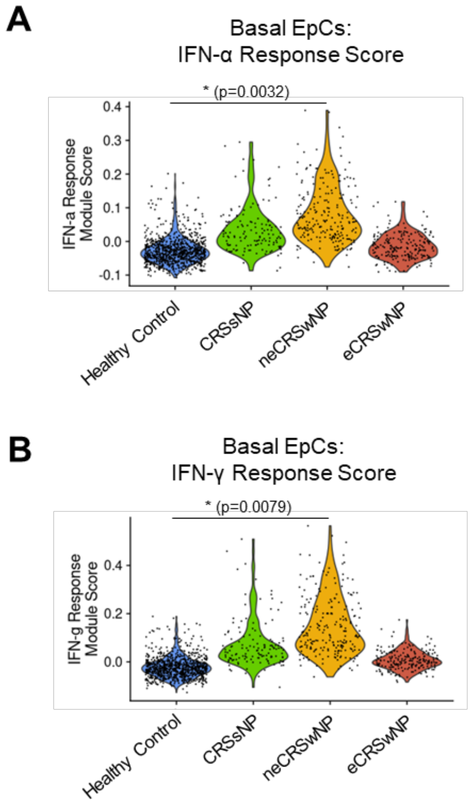


**Fig E6: Interferon response scores are elevated in neCRSwNP basal EpCs**

A) Violin plot of the IFN-α response scores in basal EpCs from the Wang scRNA-seq dataset. Each dot represents one basal EpC. The non-parametric Mann-Whitney U test was performed using the mean score for each donor (control n=5, CRSsNP n=5, neCRSwNP n=5, eCRSwNP n=6). Relevant statistical results are labeled where * denotes $p<0.05$ and *ns* denotes $p>0.05$.

B) Violin plot of the IFN-γ response scores in basal EpCs from the Wang scRNA-seq dataset.

### Fig E7

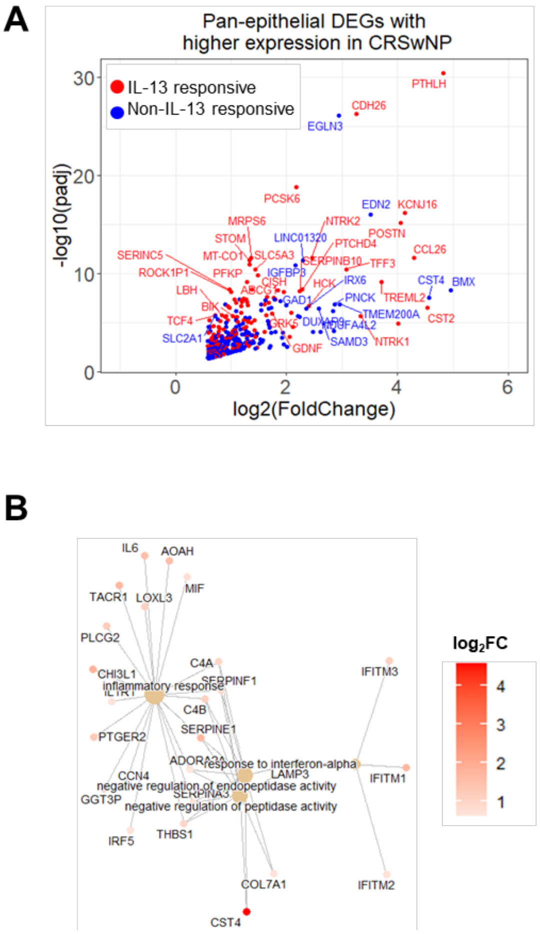


**Fig E7: Genes involved in interferon signaling are over-represented among non-IL-13-responsive pan-epithelial DEGs**

A) Volcano plot of pan-epithelial DEGs with higher expression in CRSwNP. DEGs are colored as IL-13 responsive (red) or non-IL-13 responsive (blue) based on DE testing of GSE185202.

B) Net plot of selected top over-represented gene sets (in the gene ontology biological processes database) for non-IL13-responsive pan-epithelial DEGs with higher expression in CRSwNP EpCs.

#### Fig E8

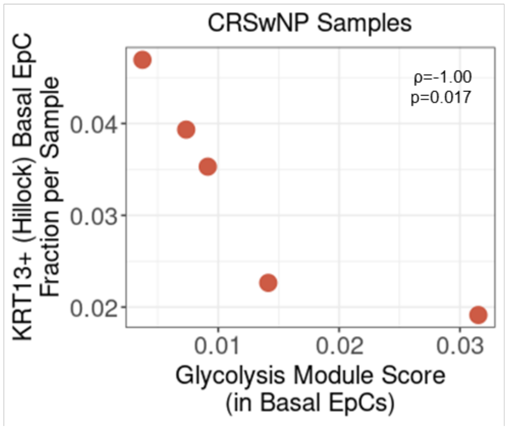


**Fig E8: EpC glycolysis negatively correlates with the fraction of KRT13+ "hillock" basal EpCs in CRSwNP**

Scatterplot of the mean glycolysis score in basal EpCs vs fraction of KRT13$^{+}$ "hillock" basal EpCs recovered by scRNA-seq from the 5 CRSwNP samples in the Kotas scRNA-seq dataset. ρ indicates Spearman's *rho*.

**Fig E9**

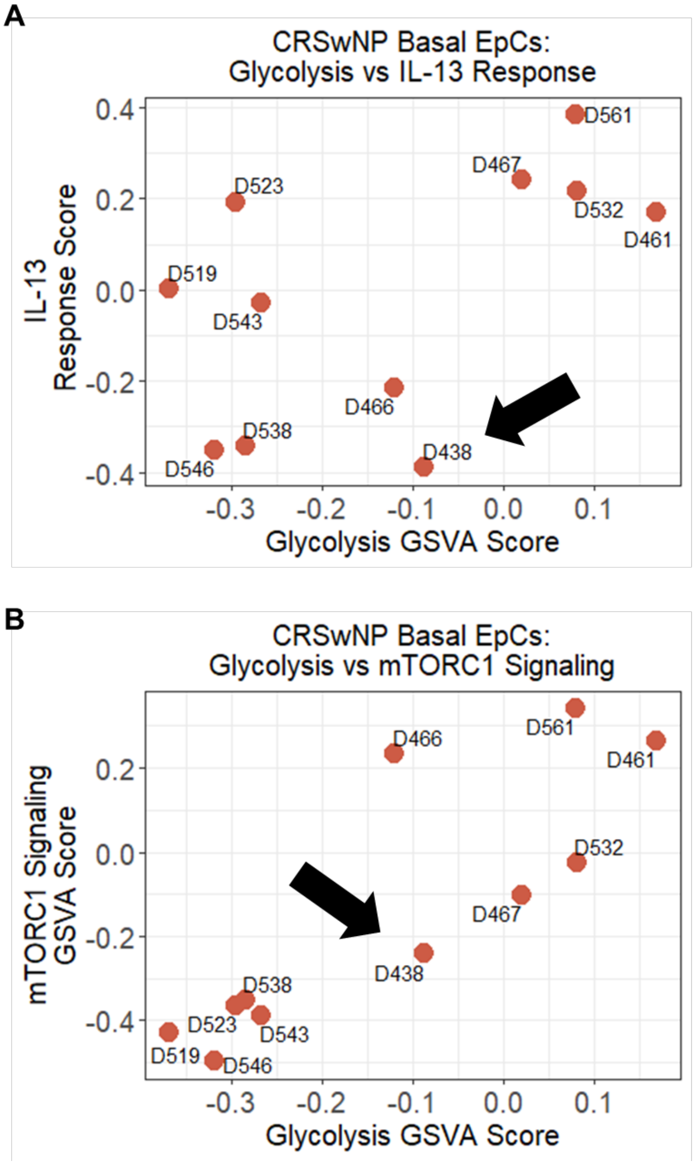


**Fig E9: Glycolysis and mTORC1 signaling were preserved despite dupilumab administration in one subject with AERD (D438)**

A) Scatterplot of glycolysis GSVA score vs IL-13 response score in CRSwNP basal EpCs. D438 (designated with the black arrow) had received dupilumab.

B) Scatterplot of glycolysis GSVA score vs mTORC1 signaling GSVA score in CRSwNP basal EpCs. D438 (designated with the black arrow) had received dupilumab.
